# Phosphorylation code of human nucleophosmin includes four cryptic sites for hierarchical binding of 14-3-3 proteins

**DOI:** 10.1101/2024.02.13.580064

**Authors:** Anna A. Kapitonova, Kristina V. Perfilova, Richard B. Cooley, Nikolai N. Sluchanko

## Abstract

Nucleophosmin (NPM1) is the 46th most abundant human protein with many functions whose dysregulation leads to various cancers. Pentameric NPM1 resides in the nucleolus but can also shuttle to the cytosol. NPM1 is regulated by multisite phosphorylation, yet molecular consequences of site-specific NPM1 phosphorylation remain elusive. Here we identify four 14-3-3 protein binding sites in NPM1 concealed within its oligomerization and α-helical C-terminal domains that are found phosphorylated *in vivo*. By combining mutagenesis, in-cell phosphorylation and PermaPhos technology for site-directed incorporation of a non-hydrolyzable phosphoserine mimic, we show how phosphorylation promotes NPM1 monomerization and partial unfolding, to recruit 14-3-3 dimers with low-micromolar affinity. Using fluorescence anisotropy we quantified pairwise interactions of all seven human 14-3-3 isoforms with four recombinant NPM1 phosphopeptides and assessed their druggability by fusicoccin. This revealed a complex hierarchy of 14-3-3 affinities toward the primary (S48, S293) and secondary (S106, S260) sites, differentially modulated by the small molecule. As three of these 14-3-3 binding phospho-sites in NPM1 reside within signal sequences, this work highlights a key mechanism of NPM1 regulation by which NPM1 phosphorylation promotes 14-3-3 binding to control nucleocytoplasmic shuttling. It also provides further evidence that phosphorylation-induced structural rearrangements of globular proteins serve to expose otherwise cryptic 14-3-3-binding sites that are important for cellular function.

## Introduction

Nucleophosmin 1 (NPM1, or B23, numatrin, NO38) is an extremely abundant human protein (top 0.25%, according to the PaxDB database [1]) with up to 30 known sites of phosphorylation that are thought to regulate its activity and interaction with other proteins. NPM1 forms stable pentamers and supramolecular assemblies inducing liquid-liquid phase separation (LLPS), and is the main component of the membraneless organelle nucleolus [2,3]. NPM1 regulates ribosome biogenesis, histone assembly, centrosome duplication, DNA integrity and cell stress responses [2,4–7]. The regulation of NPM1 functions is accompanied with its phosphorylation, oligomer disassembly and nucleocytoplasmic shuttling [7–14]. Many mutations in NPM1 cause its permanent egress to the cytoplasm and induce various cancer types including acute myeloid leukemia (AML) [15]. However, the exact mechanisms underlying NPM1 regulation, including multi-site phosphorylation and recognition by the major phosphoserine/phosphothreonine (pS/pT)-binding protein known as 14-3-3, are not well understood.

14-3-3 proteins were the first phospho-Ser/Thr binding protein modules discovered [16] and belong to the top 1% most abundant human proteins [1]. Composed primarily of α-helices, the ∼30 kDa 14-3-3 subunits assemble into homo- or heterodimers containing one groove per subunit that bind specific phospho-motifs located internally or at the C-terminus of their partner proteins [17] (Fig. 1A). As dimers, 14-3-3 proteins can bind phosphorylated clients at a single phospho-site or multi-valently through multiple phospho-sites. The complexity of 14-3-3/client interactions is evident when considering that clients commonly have more than two 14-3-3-binding motifs and can thus bind 14-3-3 in a variety of multi-valent modes depending on which combination of sites are phosphorylated, i.e. their “phosphorylation code”. In addition, humans express seven 14-3-3 subtypes encoded by separate genes, which are historically termed ‘isoforms’ and designated by the Greek letters (β (beta), γ (gamma), ε (epsilon), ζ (zeta), η (eta), σ (sigma), τ (tau)); these isoforms are highly conserved but differ by the affinity of interaction they have with their phospho-targets [18]. While the 14-3-3 interactome comprises hundreds of different proteins, only a minority have been well-studied when in complex with 14-3-3.

**Fig. 1.**
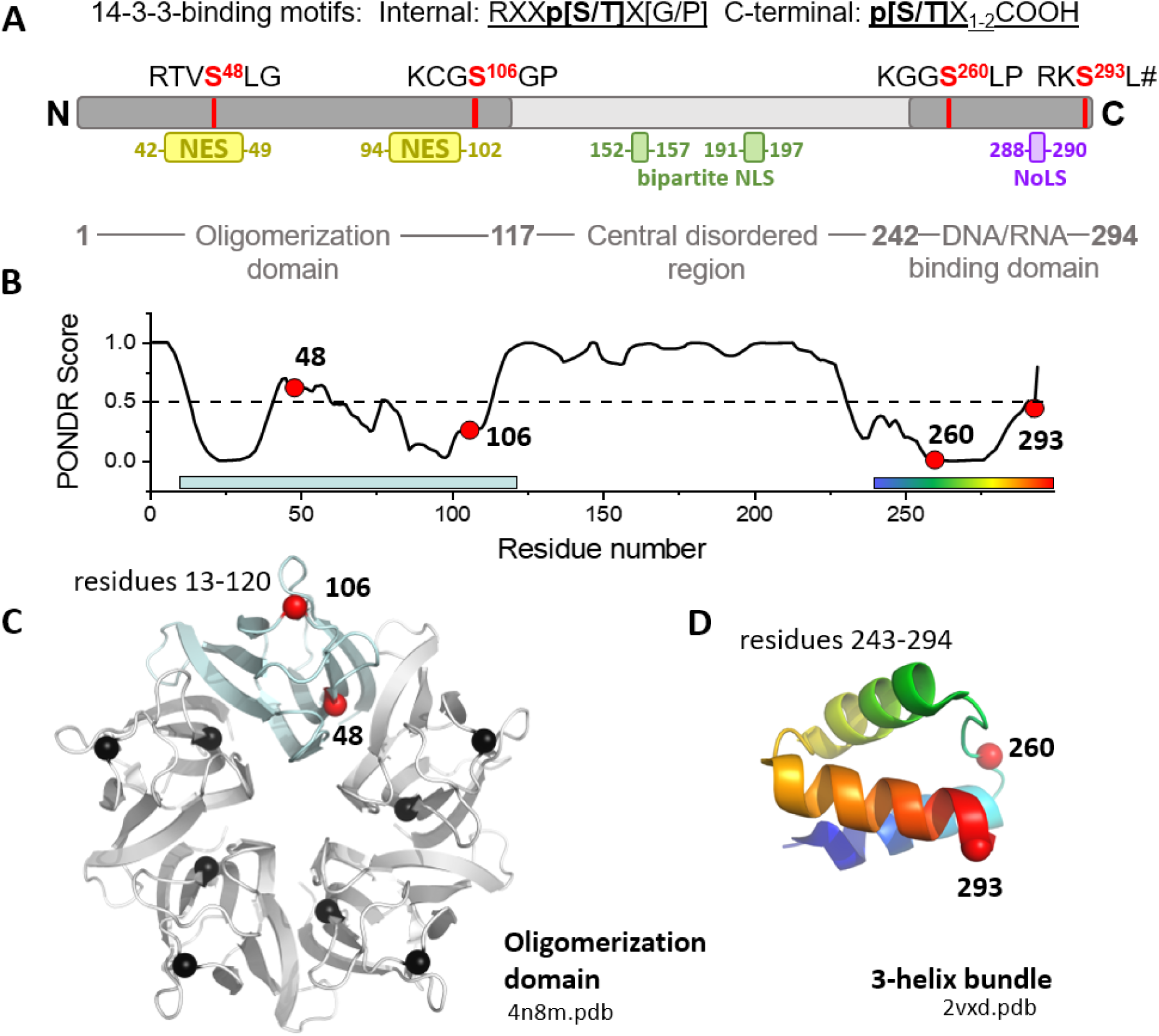
Human NPM1 protein. A. Domain structure of NPM1 showing the location of several internal and the C-terminal tentative 14-3-3-binding phosphoserines and the signal sequences (NES, NLS, NoLS). # designates the carboxyl group. B. Intrinsic disorder propensity throughout the NPM1 sequence. The two ordered regions correspond to the known oligomerization (OD) and C-terminal domain (CTD) shown on panels C and D, respectively. Red spheres indicate the location of the phosphoserines 48, 106, 260 and 293.

Recent work has implicated 14-3-3 in regulating NPM1 function. For example, centrosome amplification depended on cytosolic NPM1 cooperating with 14-3-3 [14,19] in an NPM1 phosphorylation-dependent manner [20]. Specifically, 14-3-3ε was found among the NPM1 interactors in the nucleolus [5], though another study could not reproduce the direct 14-3-3/phospho-NPM1 interaction [21]. Reasons for this discrepancy are not clear but it could be because NPM1/14-3-3 interactions depend on the specific pattern of NPM1 phosphorylation across multiple sites. Intriguingly, recent crystal structures have provided snapshots of 14-3-3 interaction with two NPM1 phospho-fragments, around pSer48 [22] and pSer293 [23]. NPM1 phosphorylation at Ser48 by prosurvival kinase Akt led to NPM1 pentamer disassembly and exit from the nucleolus [7], and so not surprisingly NPM1 phosphorylated at Ser48 is prevalent in human tumors [7]. Yet, the details of the 14-3-3/NPM1 interaction were not analyzed further and the hierarchy of these two (and potentially other) NPM1 phospho-sites in controlling complexation, remain unknown.

In this work, we addressed these questions by dissecting the interaction of 14-3-3 proteins with unphosphorylated and phosphorylated recombinant NPM1 variants. Current approaches to decipher the roles of single and multi-site phosphorylations in context of the whole protein are limited as phosphosites are not always solvent accessible for kinases, and the complexity of kinase specificity and activation mechanisms can lead to incomplete and/or off-target phosphorylation. Further, their inadvertent exposure to phosphatases leads to heterogeneity in sample preparation. Although still widely used because of the simplicity of their incorporation and stability, phospho-mimicking mutations (S/T → D/E) are rather poor mimics of pS/pT. Here, we use a combination of in-cell kinase-mediated phosphorylation strategies [24–27] as well as site-directed translational installation of a non-hydrolyzable phosphonate analog of phosphoserine [28,29] to generate an array of NPM1 phospho-variants and peptides to study their interactions with 14-3-3 proteins. We show that native NPM1 forms a stable pentamer generally resistant to phosphorylation, but once phosphorylated it disassembles into monomers that expose the phospho-sites for complexation with 14-3-3. Measuring the affinity of four putative 14-3-3 phospho-sites to all seven 14-3-3 isoforms reveals a hierarchical binding mechanism that supports the relevance of several structurally distinct, multi-valent NPM1/14-3-3 binding complexes. Additionally, we identified a selective, differential modulation of these interactions by a small molecule, fusicoccin (FSC), depending on a given NPM1 phospho-motif – from 1.4-fold inhibition to 115-fold stabilization. These results provide important clues regarding NPM1 regulation and how therapeutics targeting 14-3-3/NPM1 complexes could be used to treat disease, while also highlighting the utility of recently developed genetic code expansion (GCE) tools for generating and studying phosphorylated proteins and peptides.

## Results

### 1. Phosphorylation drives NPM1 oligomer dissociation

Human NPM1 (294 residues) is composed of an N-terminal oligomerization domain (OD), a long central unstructured region and a C-terminal nucleic acid-binding domain (CTD) featuring a C-terminal three-helical bundle [30] (Fig. 1A and B). Two minor NPM1 isoforms are shorter and lack either the stretch 195-223 in the unstructured region (isoform 2) or the C-terminal 35 residues (isoform 3). The NPM1 sequence contains over 30 documented phospho-S/T sites detected *in vivo* [31], including over 10 located in the OD (Table 1). Many of these sites represent good candidate sites for PKA phosphorylation [32], in particular Ser48 and Ser106 within the OD, and Ser260 and Ser293 within the CTD (Fig. 1A-D). At least three of these sites are located in proximity to the intracellular localization signals found within NPM1 (Fig. 1). Previous work on the OD of NPM1 has proposed that sequential phosphorylation starting from the OD periphery disassembles NPM1 oligomers and promotes their order-to-disorder transition exposing additional sites [8,9]. Yet, the effect of NPM1 phosphorylation on the structure and function of full-length NPM1 is less well understood, especially given the multiplicity of its phosphosites. Intriguingly, some of them, including Ser48, Thr78 and Ser112, are located in the subunit interfaces within the OD (Fig. 1C and 2A) and hence are likely to play regulatory roles.

**Fig. 2.**
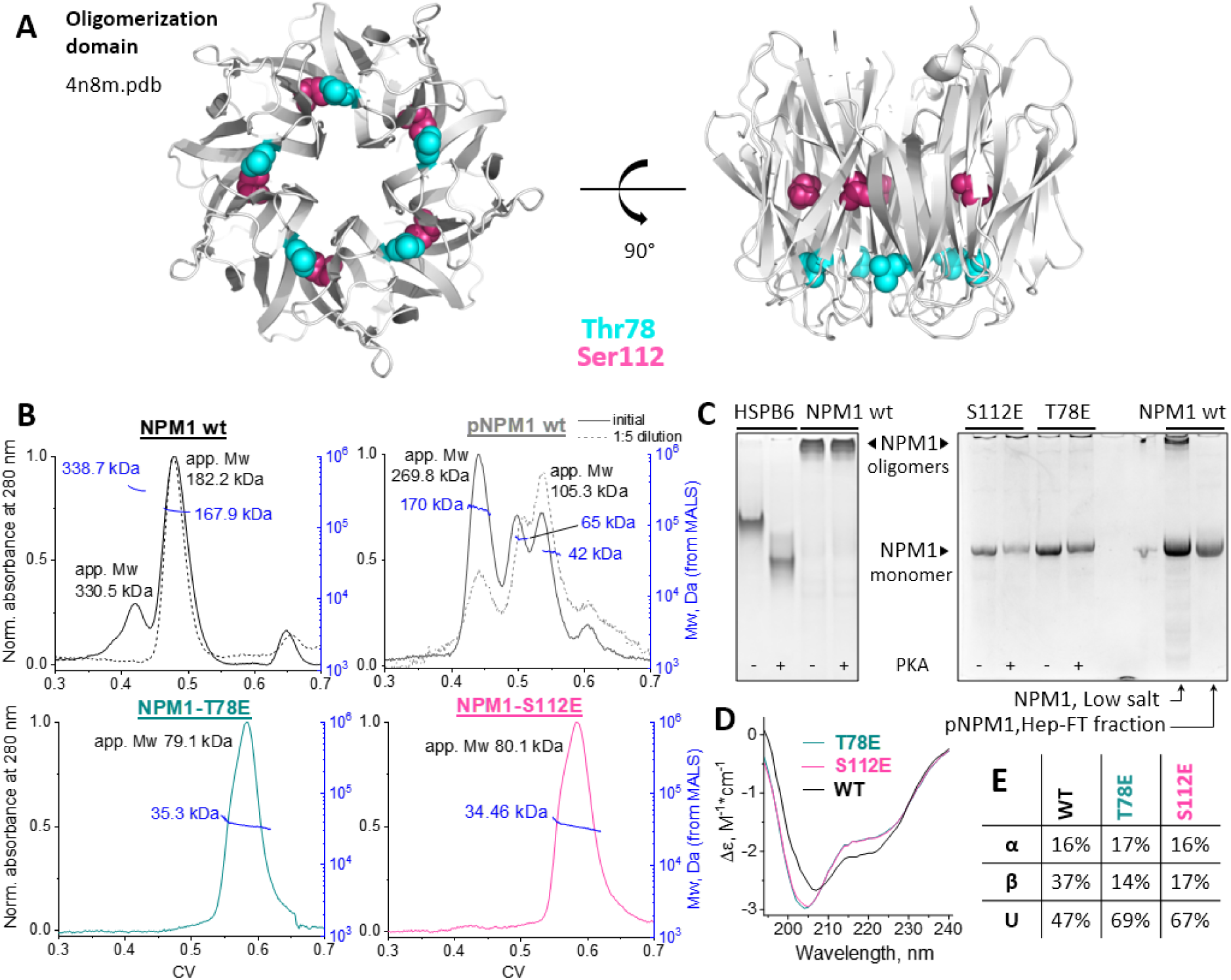
Disruption of NPM1 pentamer by phosphorylation and phosphomimicking mutations. A. NPM1 OD pentamer showing the location of the Thr78 and Ser112 residues in two orthogonal projections. B. SEC-MALS analysis of the wild-type full-size NPM1 expressed in the absence or in the presence of PKA, or of T78E and S112E mutants mimicking phosphorylation at the pentameric interface. Dashed lines correspond to the diluted sample. Absolute Mw values determined from MALS data are shown next to the Mw distributions across the selected peaks in blue. Apparent Mw values determined from column calibration with standard proteins are also indicated in black. C. Native-PAGE showing the electrophoretic mobility of NPM1 species before and after phosphorylation or dialysis against low-salt buffer. The last lane shows the mobility of the phosphorylated NPM1 fraction unable to bind to heparin. D. Far UV-CD spectra of the wild-type NPM1 or its muteins. E. Deconvolution of the CD spectra of the wild-type NPM1 and its muteins T78E and S112E by CDSSTR algorithm of the Dichroweb server [62] yielding percentages of the secondary structures: alpha-helices (α), beta-strands (β) and unstructured regions (U).

**Table 1.**
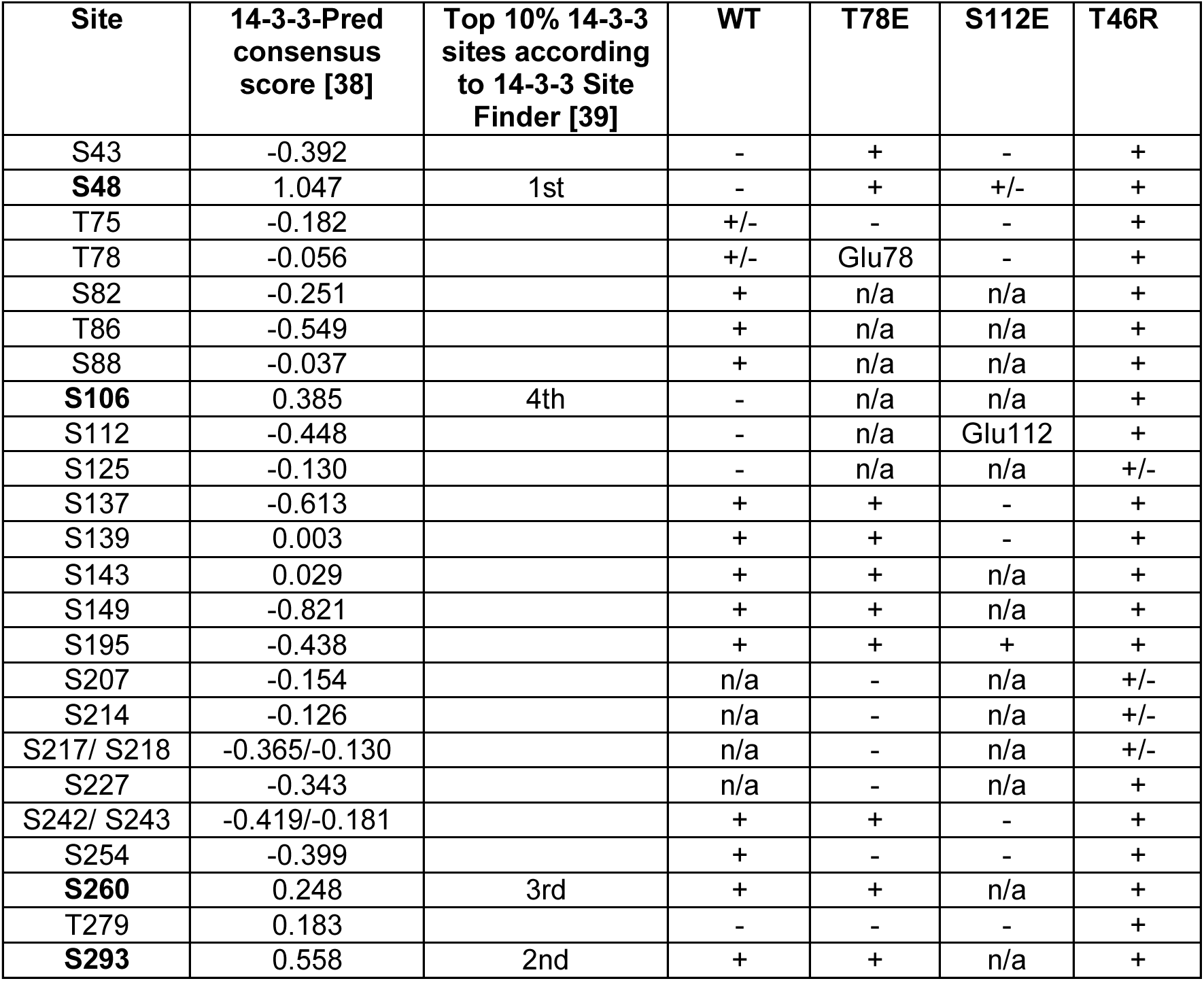
Phosphorylation of PhosphoSitePlus-listed phospho-S/T sites within human NPM1 variants upon their bacterial co-expression with PKA. Sites having Pro in position +1 relative to phosphorylatable Ser/Thr are not considered due to inability to recruit 14-3-3.

Recombinant full-length human NPM1 folds into a very stable oligomer. SEC-MALS analysis revealed a major peak with a Mw of 168 kDa (Fig. 2B), which is consistent with that of NPM1 pentamer (calculated mass 168 kDa) [6,8]. We occasionally observed a minor peak of decamer (Mw from SEC-MALS - 339 kDa), likely representing a dimer of pentamers [33]. Prolonged *in vitro* phosphorylation by PKA of these NPM1 oligomers that fail to enter the native gel did not dissociate them (Fig. 2C). To confirm PKA was functional under these conditions, a reference PKA substrate known as small heat shock protein HSPB6 [34] was indeed stoichiometrically phosphorylated (Fig. 2C). Since *in vitro* PKA-mediated phosphorylation of NPM1 was not feasible, we attempted a more effective strategy in which NPM1 phosphorylation was conducted in *E.coli* cells by co-expression with PKA, which has proven efficient for obtaining multiply phosphorylated proteins such as human Tau [24] and SARS-CoV-2 nucleoprotein [25].

Surprisingly, PKA co-expression for as long as 48 h resulted in only partial phosphorylation of NPM1, as judged from the appearance of smaller NPM1 oligomers (42-65 kDa) on SEC being in equilibrium with the NPM1 pentamer (SEC-MALS derived Mw ∼170 kDa, Fig. 2B). Interestingly, NPM1 co-expressed with PKA contained two protein subpopulations differing by the affinity for heparin, with the heparin flow through fraction containing particles with fast migration on native PAGE and a larger elution volume on SEC (Fig. 2C and Supplementary Fig. 1). This tentatively phosphorylated NPM1 population matched the native PAGE mobility of NPM1 dialyzed against low salt buffer, a treatment known to disrupt NPM1 oligomer [8,35], as well as NPM1 mutants mimicking phosphorylation of Thr78 or Ser112 within the OD (Fig. 2C). The dissociated state of T78E and S112E variants of NPM1 was confirmed by SEC-MALS analysis revealing ∼35 kDa absolute mass in each case (calculated NPM1 monomer mass is 33.6 kDa) (Fig. 2B). From these results we conclude this in-cell PKA-treated NPM1 population is primarily dissociated and monomeric. Interestingly, the T78E and S112E variants of NPM1 eluted on SEC with a higher apparent Mw than expected for a globular protein (∼80 kDa) even though MALS confirmed their monomeric status, indicating they adopt a substantially expanded/disordered conformation (Fig. 2B). Consistent with this, both phosphomimicking mutations resulted in an altered far-UV CD spectrum of NPM1 with an enhanced contribution of the unfolded regions (Fig. 2D), most likely derived from the disrupted interfaces in the OD. Indeed, secondary structure content analysis indicated that the muteins had ∼20-22% more unstructured regions at the expense of the β-strands, which reduced concomitantly by 20-23% (60-67 residues), relative to those of the wild-type NPM1 (Fig. 2E). Given the remaining β-strand contribution (∼14-17%), a complete OD unfolding accompanying NPM1 monomerization did not take place and the C-terminal helical bundle remained unaffected (16-17% of α, or 47-50 residues), in agreement with the data that it retains helical even in isolation [30,36] (Fig. 2E).

### 2. Oligomerization domain of NPM1 conceals key phosphorylation and 14-3-3-binding sites

Having obtained partially phosphorylated NPM1, we asked if such phosphorylation is sufficient to trigger NPM1 interaction with 14-3-3 proteins and applied SEC, which previously has proven to be especially effective for detecting 14-3-3 complexes [25,34,37]. As expected, the mixture of unphosphorylated NPM1 with human 14-3-3γ produced two separate peaks on the elution profile, corresponding to the individual peaks of non-interacting NPM1 oligomer and 14-3-3γ dimer (Supplementary Fig. 2). This agrees well with the previous observation and the interaction of these and many other proteins with 14-3-3 is phosphorylation-dependent [16,20]. However, under the conditions used we did not detect any significant interaction also between 14-3-3γ and NPM1 in-cell phosphorylated by PKA (Supplementary Fig. 2), suggesting that despite extensive phosphorylation and partial monomerization, NPM1 sites crucial for 14-3-3 recruitment remained masked and not phosphorylated. In favor of this assumption, we still observed the main fraction of PKA-phosphorylated NPM1 in the extended oligomerized form (increased apparent Mw), which likely disassembles only upon heavy sequential OD phosphorylation [8].

To better understand the nature of the in-cell PKA-treated phosphorylated NPM1 protein, mass-spectrometric analysis detected phosphorylation of at least five sites within the OD (namely, Thr75, Thr78, Ser82, Thr86 and Ser88, Supplementary File 1), however, the corresponding phospho-motifs have negative scores with respect to 14-3-3 binding as predicted by 14-3-3-Pred [38]. The highest ranked 14-3-3-binding phospho-motif within NPM1 according to 14-3-3-Pred (consensus score 1.047) [38] and to 14-3-3 Site Finder [39] surrounds Ser48 of the OD, which is not found in the phosphorylated state in the NPM1 sample co-expressed with PKA (Table 1). Another good candidate 14-3-3 site centered on Ser106 within the OD (consensus score 0.385) [38] also remained non-phosphorylated in our co-expression system, implying that these tentative 14-3-3-binding sites are poorly accessible for phosphorylation within the oligomerized NPM1, even though they should be sites amenable to PKA phosphorylation [32].

### 3. Mutations disrupting NPM1 oligomer promote its phosphorylation and 14-3-3 binding

We therefore asked if PKA-mediated phosphorylation of candidate 14-3-3 binding sites within the OD may be facilitated by disrupting the NPM1 oligomer using phosphomimicking mutations T78E or S112E (Fig. 2). To this end, either T78E or S112E mutein was co-expressed with PKA, purified and subjected to interaction studies with 14-3-3. As expected, unphosphorylated T78E or S112E muteins did not show any interaction with 14-3-3γ, whereas the phosphorylated counterparts did (Fig. 3), indicating PKA-mediated phosphorylation of 14-3-3-binding sites had occurred. Mass-spectrometry analysis confirmed phosphorylation of multiple sites including Ser48 in both muteins (Table 1). Therefore, we asked if phosphorylation exclusively at Ser48 can recruit 14-3-3 and so we attempted to obtain an NPM1 variant with a translationally incorporated non-hydrolyzable phosphoserine analog (phosphono-methylalanine, PMA) (p*S48) instead of Ser48 located with the putative 14-3-3-binding motif **R**TVS^48^LG. PMA encoding was made possible via the recently developed PermaPhos technology (Supplementary Fig. 3) [28]. Although the targeted NPM1 protein with PMA at Ser58 could be obtained, its yield was sufficiently low that it prevented us from complete purification and analysis of its ability to interact with 14-3-3 (Supplementary Fig. 4). The NPM1 OD contains a similar motif around Ser88 (**P**TVS^88^LG), which is phosphorylated within NPM1 but does not represent a 14-3-3-binding motif (Table 1). The p*S88 NPM1 protein could be purified with a much greater yield and indeed did not show any detectable interaction with 14-3-3γ (Supplementary Fig. 4). Both p*S48 and p*S88 NPM1 proteins with single modifications in the OD were sufficient to destabilize the NPM1 oligomer, provoking its dissociation, while the p*S48 variant was much more efficient in doing so (Supplementary Fig. 4).

**Fig. 3.**
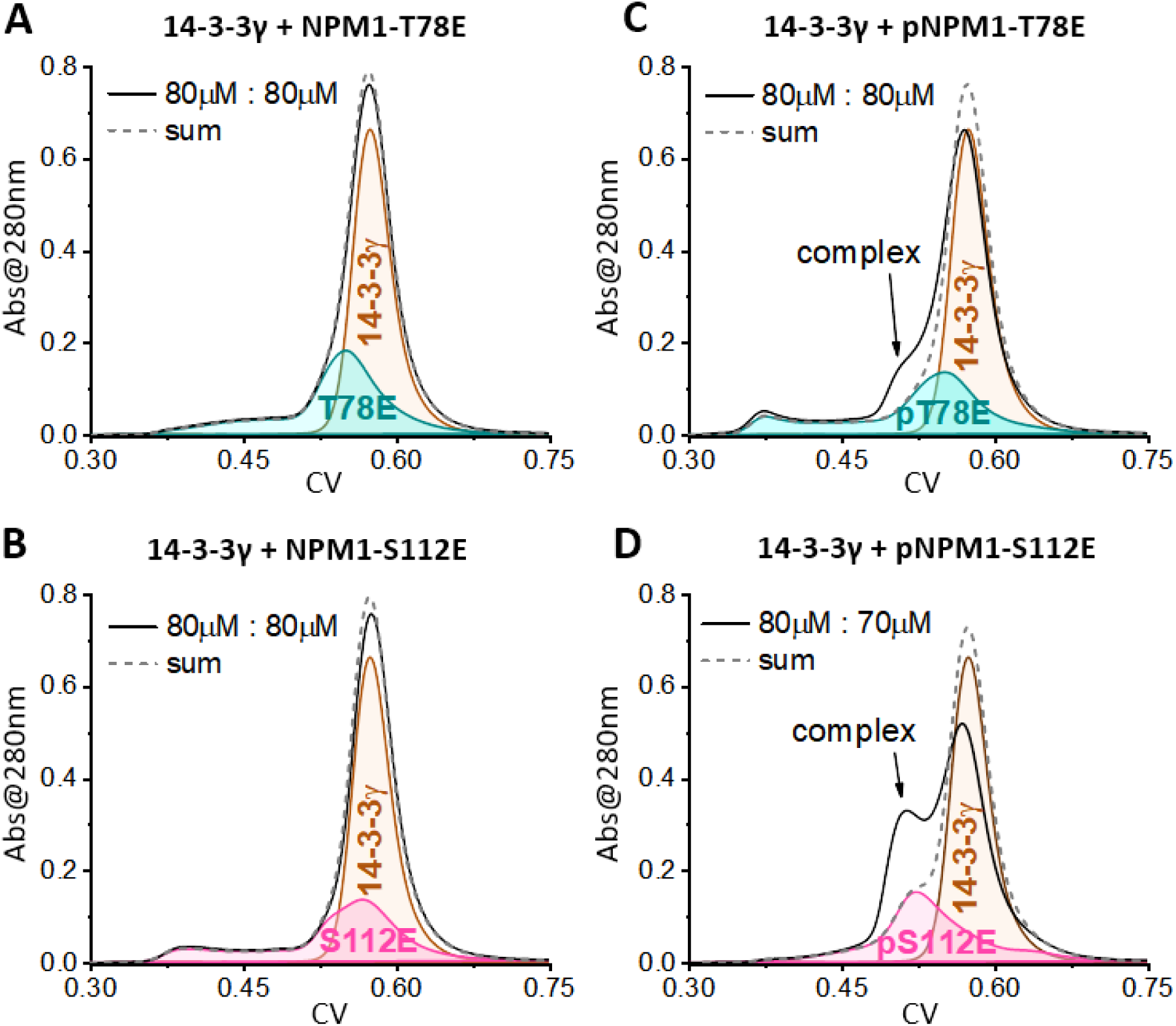
Interaction of phosphomimicking mutants of NPM1 with 14-3-3γ. A and B. SEC profiles of individual 14-3-3γ, individual unphosphorylated NPM1 T78E (A) and S112E (B) mutants or the mixtures of 14-3-3γ with either mutein. C and D. SEC profiles similar to those presented in A and B but with NPM1 muteins in-cell phosphorylated by PKA. Complex formation with 14-3-3γ is observed only in the case of phosphorylated NPM1 muteins. Column: Superdex 200 Increase 5/150 (flow rate 0.45 ml/min).

The Ser48-containing motif is the best 14-3-3-binding candidate in human NPM1 (Table 1); its phosphorylation was shown to trigger 14-3-3 binding to the surrounding peptide fragment [22]. T78E, and especially S112E, NPM1 versions containing partially phosphorylated Ser48 do recruit 14-3-3γ, but still the complex formation was limited by phosphorylation efficiency (Fig. 3). In order to improve the efficiency of Ser48 phosphorylation and thereby ensure maximum potential for NPM1 binding to 14-3-3, we modified the Ser48-motif by making it closer to the PKA and 14-3-3 consensus motifs via a T46R amino acid replacement. The expected effects were threefold. First, it would block undesired phosphorylation of Thr46, which would be unfavorable for 14-3-3 binding at Ser48 [40] and sometimes was detected by mass-spectrometry of NPM1 phosphoforms (Supplementary File 1). Second, it would make a R**R**VSLG motif better phosphorylated by PKA (R**R**XS consensus [41]) and recognized by 14-3-3 (R**R**XpSXG/P consensus [42]). Third, given its location in NPM1 structure (Fig. 4A), the T46R replacement would destabilize intersubunit contacts within the NPM1 pentamer, tentatively promoting its dissociation and unmasking sites for phosphorylation and 14-3-3 binding without additional mutations/modifications (e.g. T78E and S112E).

**Fig. 4.**
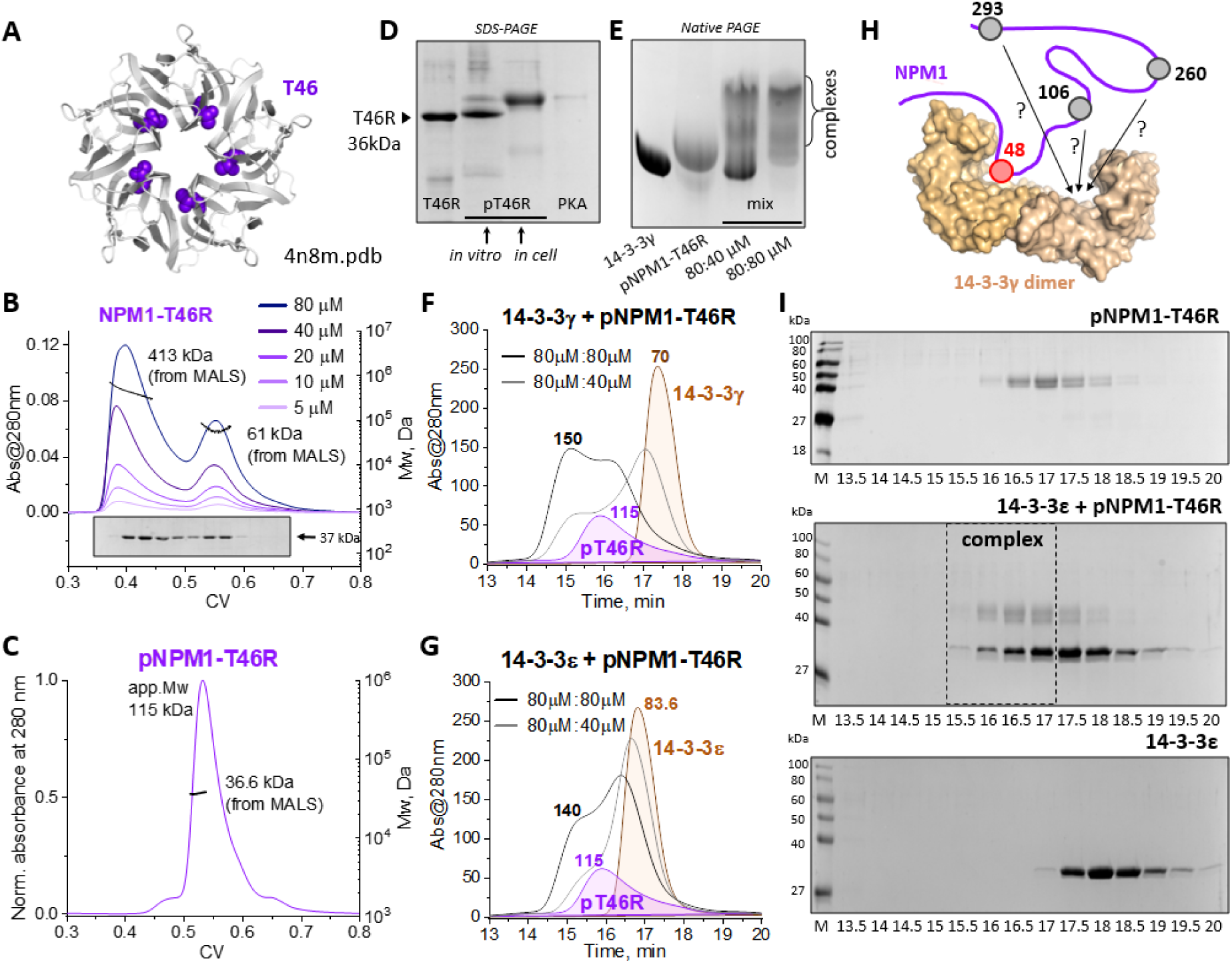
Engineered phosphorylated NPM1 monomer is capable of tight interaction with 14-3-3γ and 14-3-3ε. A. NPM1 OD structure showing the location of Thr46. B. SEC profiles showing that the T46R mutation in the full-size NPM1 induces pentamer disruption and concentration-dependent self-association via alternative interfaces. Mw distribution derived from the MALS data for the highest T46R concentration is shown for the two peaks. C. SEC-MALS profile showing that the in-cell phosphorylated T46R mutant of the full-size NPM1 forms an expanded monomer. Load concentration is 40 μM. D. T46R phosphorylation by PKA in cells, but not *in vitro*, leads to an upward shift of the electrophoretic mobility on SDS-PAGE. E. Phosphorylated full-size T46R mutein forms complexes with 14-3-3γ visualized by native-PAGE. Two molar ratios 14-3-3:NPM1 (μM) were analyzed. F and G. Phosphorylated T46R mutein is capable of forming stable complexes with 14-3-3γ (F) and 14-3-3ε (G). Note a much less efficient interaction in the case of 14-3-3ε. Apparent Mw values determined from column calibration with standard proteins are indicated above the peaks. Two molar ratios were used for each 14-3-3. H. Schematic model of the 14-3-3 dimer interacting with the phosphorylated NPM1 monomer via pS48 and some other yet undefined site(s). I. SDS-PAGE analysis of the fractions collected during SEC runs presented on panel G for individual phosphorylated T46R (top), 14-3-3ε (bottom) or their mixture (middle). Complex fractions are outlined by a dashed rectangle. Analogous data for 14-3-3γ are found in Supplementary Fig. 5.

The unphosphorylated purified T46R mutein displayed two peaks on the SEC profile (SEC-MALS-derived Mw values 60 and 410 kDa) and they interconverted based on protein concentration (Fig. 4B). We did not observe a stably monomeric T46R form, perhaps because disturbing the native subunit interfaces in the pentamer caused formation of another interface that stabilized the protein in an altered dimer and a higher-order oligomer. The phosphorylated T46R mutein (obtained via in-cell PKA phosphorylation) exhibited a single SEC peak with the absolute Mw close to the NPM1 monomer (36.6 kDa vs 33.6 kDa; Fig. 4C). Yet its increased apparent Mw of ∼115 kDa, based on its SEC elution properties, indicated that it is even more extended than the unphosphorylated T78E or S112E muteins, likely because of multisite phosphorylation (Fig. 2B). This multiply phosphorylated pNPM1-T46R form produced via in-cell PKA phosphorylation displayed slower electrophoretic mobility on SDS-PAGE gels indicative of heavy phosphorylation. And consistent with the notion that in-cell phosphorylation is more efficient, we could not reproduce this form via *in vitro* PKA phosphorylation (Fig. 4D). Indeed, mass-spectrometry confirmed multisite phosphorylation (Supplementary File 1) including at all four candidate 14-3-3-binding sites pS48, pS106, pS260 and pS293 (Table 1). Mixing pNPM1-T46R with 14-3-3γ resulted in an appearance of at least two bands corresponding to the protein complex, one of which was observed only when NPM1 was added to 14-3-3γ in substoichiometric quantities suggesting a stepwise complex assembly mechanism (Fig. 4E). This dose-dependent 14-3-3γ/pNPM1-T46R interaction was further confirmed on SEC, where a distinct peak of the complex appeared at substoichiometric NPM1 concentrations (Fig. 4F). A similar complex was observed for another 14-3-3 isoform, 14-3-3ε, although the relative abundance of the 14-3-3ε/pNPM1-T46R complex peak was substantially lower (Fig. 4F and G), in agreement with the earlier reported differences in binding efficiency these isoforms display in phosphotarget complex formation [18,25,26]. The formation of the complex between the 14-3-3 dimer and hyperphosphorylated NPM1 (Fig. 4H) was fully confirmed by SDS-PAGE of fractions collected during the SEC runs (Fig. 4I and Supplementary Fig. 5).

The much improved efficiency of 14-3-3 binding to the phosphorylated T46R mutein indicated the significant role for Ser48 phosphorylation in 14-3-3 recruitment, although the presence of all four candidate 14-3-3 binding phospho-motifs (pS48, pS106, pS260 and pS293) in this NPM1 enabled simultaneous binding of different pairs of phosphosites (Fig. 4H), all satisfying the apparent stoichiometry of one 14-3-3 dimer per one NPM1 monomer as suggested by the SEC experiment (Fig. 4F and G).

### 4. 14-3-3 affinity hierarchy of NPM1 phospho-motifs

To disentangle the hierarchy of 14-3-3 interaction with each of the four tentative 14-3-3-binding phosphosites, we set out to quantify these interactions using NPM1 peptide fragments (Fig. 5A). The S48, S106 and S260 peptides corresponding to the internal 14-3-3-binding motifs [42] (Fig. 1A) were recombinantly obtained as His_6_-bdSUMO-peptide-YFP fusions (Fig. 5B) cleavable by SUMO-hydrolase (bdSENP) and thrombin to remove the His_6_-bdSUMO and YFP tags, respectively. TAG (amber) stop codons were placed at the phosphorylatable sites 48, 106 or 260 in these peptides to direct the translational incorporation of p*S using the PermaPhos technology [28]. This approach secured the stoichiometry of phosphorylation by blocking dephosphorylation that could occur during protein expression and purification, while also ensuring the efficient recognition by 14-3-3 proteins that is almost indistinguishable from that of phosphoserine [28,43]. Accumulation of truncated products prematurely terminated at TAG codons could be neglected since these are non-fluorescent and significantly shorter due to the placement of the YFP tag C-terminal to the TAG codon. The YFP tag also helped during modification of the peptide’s N terminus by FITC and removal of the label excess, and was cleaved off later to release the corresponding FITC-labeled peptides containing p*S at the desired position (See Methods). Since this translational encoding of p*S approach with YFP could not be used for the C-terminal 14-3-3 motif around the penultimate Ser293 residue (which must be followed by the native Leu residue containing a free carboxyl-group), the corresponding peptide was produced as a His_6_-bdSUMO-fusion co-expressed with PKA (See Methods and Fig. 5B).

**Fig. 5.**
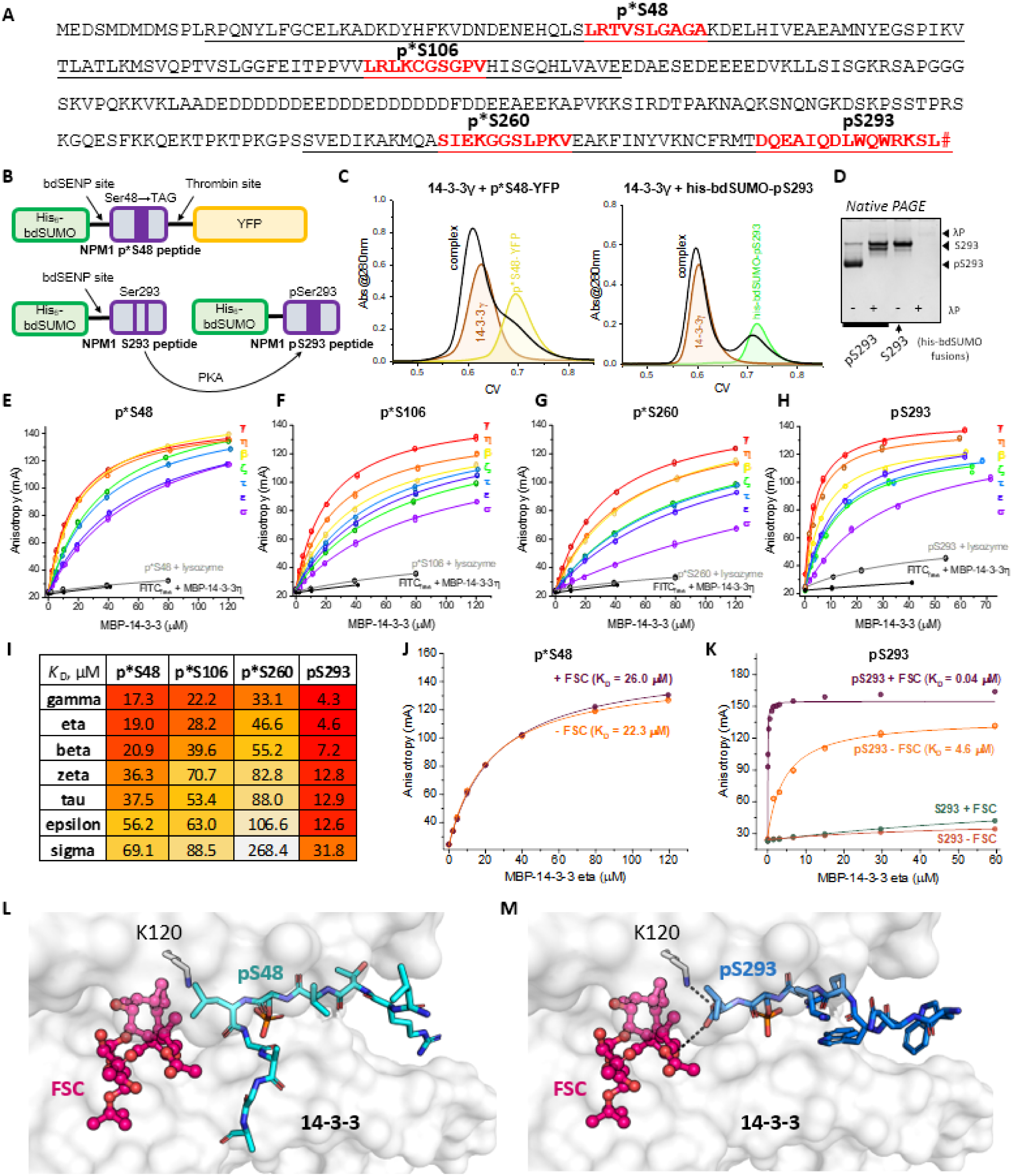
Interaction of seven human 14-3-3 isoforms with four NPM1 phospho-motifs. A. Amino acid sequence of NPM1 showing the location of the 14-3-3-binding motifs (red) within the ordered NPM1 domains (underlined). # designates the COOH group. p*S designates that the central phosphorylated residue was replaced by the non-hydrolyzable Ser analog PMA (three internal motifs). In the C-terminal motif, a canonical phosphoserine was formed by in-cell phosphorylation by PKA. Both peptide types were obtained in *E. coli* as fusion proteins (B) (see Methods). C. SEC profiles showing the interaction of 14-3-3γ with the NPM1 phosphopeptides in the context of fusion proteins with YFP or his-bdSUMO. D. Native PAGE showing the phosphorylation status of the his-bdSUMO-tagged S293 peptide co-expressed with PKA in *E.coli*. λP designates lambda phosphatase used for dephosphorylation. E-H. Binding curves obtained throughout the titration of the FITC-labeled NPM1 phosphopeptides by increasing amounts of MBP-14-3-3 isoforms. Control curves correspond to titration of free FITC by MBP-14-3-3 or of FITC-peptide by lysozyme. I. Equilibrium dissociation constants (*K*_D_) obtained by fluorescence anisotropy and characterizing the affinities of the four FITC-labeled NPM1 phosphopeptides to the seven human 14-3-3 isoforms. The values are accompanied by a color gradient for convenience. J-K. The effect of fusicoccin (FSC) on the interaction of MBP-14-3-3η with the FITC-labeled p*S48 and pS293 peptides. For pS293 peptide, titrations in the absence or in the presence of FSC were performed also for the unphosphorylated peptide version. The binding curves were approximated by the quadratic equation [63]. L. Structural model showing the absence of steric clashes and stabilizing interactions between FSC and the pS48 phosphopeptide based on the 14-3-3ζ/NPM1-pS48 peptide structure (PDB 8ah2 [22]). M. Structural model showing the canonical FSC-stabilized interaction between 14-3-3σ and the pS293 phosphopeptide (PDB 7obg and 7obh [23]) with the contacts formed between the peptide’s COOH group and 14-3-3 and FSC.

First, all NPM1 fragments displayed the interaction with 14-3-3γ as YFP or His_6_-bdSUMO fusions on SEC, while the unphosphorylated S293-peptide taken as a negative control showed no interaction (Fig. 5C,D and Supplementary Fig. 6). Second, as isolated FITC-labeled fragments, all four NPM1 peptides showed dose-dependent and saturable interaction with each of the seven human 14-3-3 isoforms, allowing us to determine equilibrium *K*_D_ values for each protein-peptide pair (Fig. 5E-I). Of note, control experiments showed unambiguously that no interaction of 14-3-3 proteins with free FITC occurred, nor any significant interaction of the peptide fragments with noncognate proteins (MBP or lysozyme); likewise, no interaction with 14-3-3 was observed for the unphosphorylated S293-peptide (Supplementary Fig. 8). As a result, 28 physiologically relevant dissociation constants could be determined (Fig. 5I and Supplementary Fig. 8), covering a 62-fold difference between the strongest (14-3-3γ/pS293) and the weakest binding pair (14-3-3σ/p*S260). While all *K*_D_ values lied in the micromolar range, 14-3-3 isoforms exhibited the remarkable selectivity toward the NPM1 phospho-motifs, recognizing most strongly pS293, followed by p*S48, p*S106 and p*S260 peptides (Fig. 5E-I). It is unlikely that the tighter pS293 peptide affinity was the result of it having authentic phosphoserine compared to the other peptides that instead had the phosphonate derivative of phosphoserine (p*S), since prior work has shown the same peptides/proteins containing pS and p*S bind 14-3-3 similarly [28,43]. In addition, 14-3-3 does not use the bridging γ-oxygen pf pSer as a hydrogen bond acceptor, which is replaced with CH2 in p*S [28]. Further, the substantially different affinity each of the four NPM1 peptides displayed toward each of the 14-3-3 isoforms differed accordingly and in perfect agreement with larger scale interactome data [18], with gamma/eta being the strongest and sigma being the weakest phosphotarget binders.

The druggability of the 14-3-3/NPM1 phospho-motif interaction could be probed by performing titrations in the presence of the well-known stabilizer of 14-3-3 complexes, fusicoccin (FSC) (Fig. 5J,K and Supplementary Fig. 8) [23,44]. This revealed a remarkable modulation in binding properties: while we observed an insignificant inhibitory effect of FSC in the case of p*S48 NPM1 or p*S106 peptide (Fig. 5J and Supplementary Fig. 8) and a 1.4-fold inhibition in the case of the p*S260 NPM1 peptide (Supplementary Fig. 8), there was a drastic increase in affinity in the case of the C-terminal phospho-motif pS293 (Fig. 5K and Supplementary Fig. 8). In this case, we observed a 115-fold enhancement in binding for the strong phosphotarget binder, 14-3-3η, and a 71-fold enhancement for the weak binder, 14-3-3σ (Supplementary Fig. 8). Of note, FSC could not invoke the interaction of 14-3-3 with the unphosphorylated S293-peptide (Fig. 5K). The pronounced selectivity of FSC action on NPM1 phosphopeptides could be rationalized based on the available structural data. The 14-3-3-bound conformation of the pS48-phosphopeptide leaves sufficient room for the FSC molecule to simultaneously occupy its binding site in the amphipathic 14-3-3 groove but with no apparent stabilizing interactions (Fig. 5L), consistent with no detectable effect of FSC on the 14-3-3/p*S48 peptide interaction (Fig. 5J). By contrast, the structure of the ternary complex between 14-3-3σ, pS293-phosphopeptide and FSC (PDB ID 7obh [23]), reported very recently, revealed the canonical interaction of the Leu294 carboxyl group with the conserved Lys120 residue of 14-3-3 and with FSC (Fig. 5M), typical of many C-terminal phospho-motifs of 14-3-3 [23]. In this case, FSC acts to bridge interactions between the pS293-phosphopeptide and 14-3-3, thus contributing to the enhanced stabilization of the complex.

## Discussion

NPM1 is a phosphoprotein containing over 30 serines/threonines phosphorylatable *in vivo* [31]. A complete picture of the NPM1 regulation by phosphorylation remains elusive in part due to the lack of knowledge on the effect of site-specific NPM1 phosphorylation, which reflects a more general challenge associated other multisite phosphorylated proteins, including human Tau [24,45,46], CFTR [47], LRRK2 [48] and SARS-CoV-2 nucleoprotein [25,49], all of which represent established multi-site binders of 14-3-3 proteins. Being an intrinsically bivalent dimer, 14-3-3 can bind to different pairs of phosphosites within a multi-phosphorylated target protein, and it is expected that each binding mode would produce different functional or signaling outcomes depending on the location and surrounding context of the corresponding phospho-motifs [50,51]. The mechanisms that control the prevalence of each binding mode at a given time in the cell, and how they interconvert with each other for each client are poorly understood and difficult to predict, as different 14-3-3-binding phosphosites with even similar sequences can have astonishingly different affinities to a given 14-3-3 isoform [18,48]. Further, the complex mechanisms controlling the phosphorylation code, which is determined by hundreds of protein kinases and phosphatases each of which themselves are activated by distinct stimuli, make it difficult to even know which phospho-forms of a given protein are even present under a given condition. Moreover, human 14-3-3 isoforms display a hierarchy of binding affinities to a given phosphopeptide, forming roughly an affinity trend with a ∼12-fold affinity difference for the weakest binding 14-3-3 isoform, sigma, and the strongest binding 14-3-3 isoform, gamma [18]. The recent interactomics study suggests that the phosphopeptide affinities to 14-3-3 isoforms can cover as large as a 40,000-fold range, from low nM to mM [18]. Hence, multiply phosphorylated protein partners of 14-3-3 represent a great challenge and require high-throughput approaches for accurate *K*_D_ determination between 14-3-3 proteins and the corresponding phospho-fragments [52]. Importantly, such a fragmentomics approach is meaningful only once the phosphorylation of the sites in question is confirmed in context of a full-size protein, for which the phosphorylation-dependent interaction with 14-3-3 would need to be confirmed as well. For multiply phosphorylated 14-3-3 targets this task is especially challenging.

Further adding to the complexity of these challenges is 14-3-3-binding proteins can be polymorphic that undergo structural and/or oligomeric state transitions throughout their life cycle in order to controllably conceal and expose 14-3-3-binding motifs. In these cases, structurally ordered regions in the unphosphorylated state are inaccessible to 14-3-3, while phosphorylation drives necessary structural rearrangements that favor unfolding and/or dismantle oligomers. This makes such cryptic phosphorylatable motifs conditional 14-3-3-binding sites [26,53] as manifested in human STARD1, a protein known to undergo unfolding during mitochondrial importation [26].

Protein translocation between cellular compartments is often accompanied by structural transition, posttranslational modifications and interaction with other cell factors. For example, human Forkhead transcription factor FKHRL1 is phosphorylated in the nucleus by Akt kinase in response to growth/survival factor signaling and is translocated into the cytosol, where it is sequestered by binding to 14-3-3 [54,55]. 14-3-3 normally transits to and from the nucleus while complexed with other proteins to control their nucleocytoplasmic shuttling [54]. In this process, the intercompartment shuttling of 14-3-3 complexes relies on the masking or exposure of corresponding NES/NLS signals within 14-3-3 clients rather than on any such signals in 14-3-3 [54].

Here, we show that two NES signal sequences (overlapping with Ser48 and Ser106 sites) and one NoLS signal sequence (surrounding Ser293) in NPM1 are *bona fide* 14-3-3-binding phosphosites (Fig. 1). This strongly implies that the corresponding phosphorylations play a regulatory role in redirecting NPM1 between the compartments. Of note, among the three alternatively spliced NPM1 isoforms, only isoform 1 (294 residues) and isoform 2 (265 residues) contain all four sites (Ser48, Ser106, Ser260 and Ser293) for 14-3-3 binding, whereas isoform 3, a resident of the nucleus expressed in low quantities, lacks the last 35 residues on the C terminus (residues 258-294) and hence contains only Ser48 and Ser106 sites for 14-3-3 recruitment (Fig. 1). One or more C-terminal 14-3-3-binding sites appear to be missing also in deletion NPM1 mutants associated with the AML [30,56]. Therefore, such mutants cannot interact with 14-3-3 using the corresponding C-terminal NPM1 motifs. The common presence of Ser48 and Ser106 in the NPM1 isoforms and AML mutants suggests their universal importance for 14-3-3 recruitment. While these sites are located within the pentameric N-terminal domain of NPM1, like the Ser260 and Ser293 sites located within the helical C-terminal domain, all of them can be exposed upon NPM1 phosphorylation, which destabilizes their ordered structures due to the direct interference with the protein interfaces (Fig. 6). As proposed in [8,9,32,35], disassembly of the NPM1 oligomer likely occurs upon sequential phosphorylation of several sites within its oligomerization domain, starting from the most accessible ones.

**Fig. 6.**
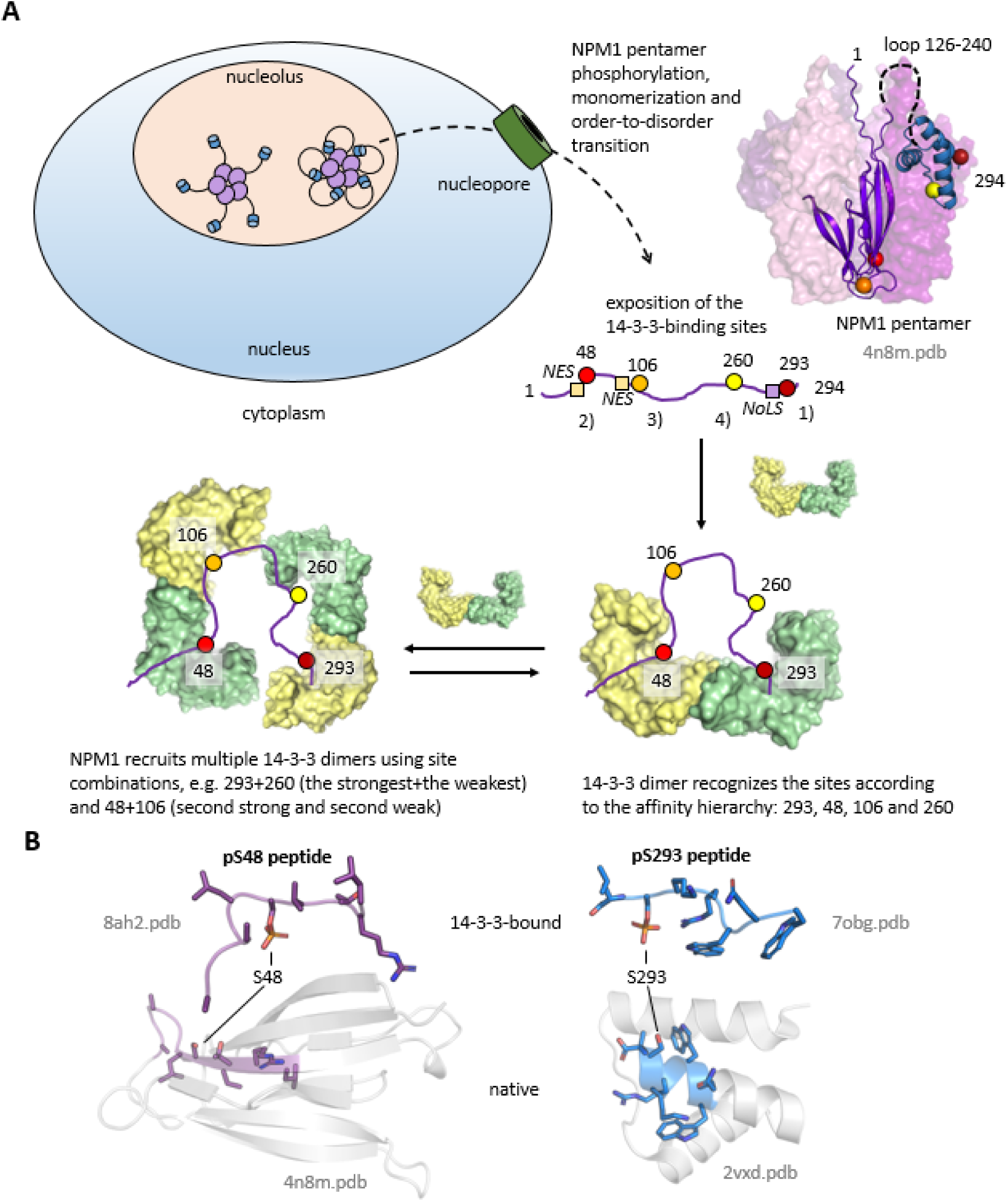
Sequestration of phospho-NPM1 by 14-3-3 in the cytoplasm via the hierarchical binding to NPM1 phospho-motifs. A. Schematic illustrating that the nucleolar NPM1 pentamer gets translocated to the cytoplasm upon phosphorylation and partial unfolding, which exposes the 14-3-3-binding sites (shown by colored circles). The sites display a pronounced 14-3-3 affinity hierarchy from strong to weak: 293, 48, 106, 260 (color-coded and numbered 1-4). Of note, 48 and 106 sites are adjacent (106) or overlap (48) with the functional NES signals, whereas 293 site is adjacent to the NoLS signal of NPM1. 14-3-3 dimer binding modes are shown for the situation of 1 or 2 dimers bound to NPM1 using the hierarchized sites. B. Structural comparison showing the conformational difference between the 14-3-3-bound peptide and the corresponding peptide fragment within native NPM1 for the pS48 (left) and pS293 (right) peptides of NPM1. Ordered structures comprising these fragments within the native NPM1 are shown by white semitransparent cartoons.

Considering that phosphorylated, dissociated NPM1 loses its ability to bind to the heparin column (a proxy of its binding to polyanionic nucleic acids) (Fig. 2), we propose that phosphorylation of nucleolar NPM1 releases NPM1-bound nucleic acids and leads to phospho-NPM1 monomerization, partial unfolding (Fig. 2) and 14-3-3 recruitment (Fig. 6). The four cryptic phosphosites occurring *in vivo* (Ser48, Ser106, Ser260 and Ser293) (Table 1) strongly differ by the affinity to 14-3-3, each showing a distinct affinity trend to 14-3-3 isoforms [18]. The hierarchy of affinities to 14-3-3 imply two main 14-3-3-binding modes: one in which the 14-3-3 dimer is bound to one NPM1 via the two strongest interaction phospho-sites (pSer48+pSer293) and a second in which two 14-3-3 dimers are recruited to NPM1 molecule via one strong and one weak phospho-site (i.e. pSer48+pSer106 and pSer260+pSer293) (Fig. 6), which would depend on local 14-3-3 abundance. The fact that the canonical and rather extended 14-3-3-bound conformations of NPM1 phosphopeptides around Ser48 and Ser293 differ from those adopted by the same peptides within native NPM1 (Fig. 6) [9,32] allows us to consider these 14-3-3-binding sites as conditional and speculate that this phenomenon is more widespread within the 14-3-3 interactome than currently recognized [26]. It is also tantalizing to suggest that 14-3-3/phospho-NPM1 interaction dissected in the present work may have significant implications in the liquid-liquid phase separation properties of NPM1, an activity relevant for maintaining stability and integrity of the membraneless organelle nucleolus [57].

The high affinity of Ser48 and Ser293 peptides to 14-3-3 (Fig. 5) is further enhanced in full-length NPM1 by the secondary sites Ser106 and Ser260, which are rather low-affinity sites on their own, because the cooperative, bidentate binding to 14-3-3 can be an order of magnitude stronger due to the avidity effects and increased local concentration of 14-3-3 sites [34,42,58]. Such low-micromolar or even nanomolar affinity would imply that phosphorylated NPM1 is efficiently sequestered by 14-3-3 in the cytoplasm upon phosphorylation (Fig. 6). NPM1 is expressed at extremely high levels in human cells and is the 46th most abundant human protein (among 19,338) with an abundance of 2605 ppm [1], which translates into ∼10 μM intracellular concentration, if the total protein concentration is taken as 3-5 mM [59]. This means that complete phosphorylation of 14-3-3-binding sites in NPM1 makes it an efficient competitor for 14-3-3 binding and a potential displacer of many low abundant and low-affinity 14-3-3 phosphotargets. Given that NPM1 phosphorylation at multiple sites occurs during mitosis [60], we propose that the interaction of the phospho-NPM1 with 14-3-3 protein hubs can systemically rewire their interactome throughout the cell cycle.

Lastly, the contrasting effect of FSC on 14-3-3 binding to NPM1 phospho-motifs from slight inhibition (Ser260) over neutral (Ser48 and Ser106) to drastic stabilization (Ser293) indicates the possibility of the site-selective druggability of the NPM1/14-3-3 interaction, which can be considered in the future chemical biology studies for combating NPM1-associated cancers. Similarly, a differential FSC action on the 14-3-3-client interactions has recently been reported on a systemic level upon neurite outgrowth stimulation [61].

## Methods

Chemicals, Molecular cloning, Protein expression and purification, Recombinant NPM1 phosphopeptides, Far-UV circular dichroism spectroscopy, In vitro phosphorylation/dephosphorylation and gel-electrophoresis, Size-exclusion chromatography, SEC-MALS, FITC labeling of recombinant NPM1 peptides, Fluorescence anisotropy and Identification of NPM1 phosphosites using LC-MS analysis are described in detail in Online Methods.

## Data availability

All data associated with the study are available upon reasonable request.

## Acknowledgements

The authors thank Prof. Tapas K. Kundu for providing the NPM1 plasmid, Prof. Gilles Travé for sharing MBP-14-3-3 plasmids, Sergey Y. Filkin for help with peptide concentration using speedvac, Andrey Tsedilin for help with LC-MS, Prof. Alexey Babakov for the provided fusicoccin preparations and Dr. Yaroslav Faletrov for fusicoccin identity verification. The study was partly supported by the Ministry of Science and Higher Education of the Russian Federation (075-15-2021-1354). 14-3-3 proteins were purified within the framework of the Russian Science Foundation grant 19-74-10031 (to N.N.S.). SEC-MALS, MALDI, LC-MS and CD measurements were done at the Shared-Access Equipment Centre “Industrial Biotechnology’’ of the Federal Research Center “Fundamentals of Biotechnology” of the Russian Academy of Sciences. This work also was aided by the GCE4All Biomedical Technology Optimization and Dissemination Center supported by National Institute of General Medical Sciences grant RM1-GM144227.

## Author contributions

N.N.S. conceived and supervised studies, A.A.K., K.V.T. and N.N.S. produced proteins and performed experiments, A.A.K, K.V.T, N.N.S. and R.B.C. analyzed data, N.N.S. wrote the paper with input from all authors.

## Supplementary data

### Online methods

#### Chemicals

All regular chemicals were of the highest purity and quality available. All water solutions in the study were prepared on the milliQ-quality water (≥18.3 MΩ/cm) and were filtered through the 0.22 µm Millipore filter system before use. Isopropyl-β-thiogalactoside (IPTG) was from Thermo Scientific (#R0392), fluorescein isothiocyanate (FITC) was from Sigma-Aldrich (#F7250-1G), thrombin was from Tehnologia-standart (#017).

#### Molecular cloning

The NPM1 gene coding for Uniprot ID P06748 protein sequence carrying uncleavable C-terminal His_6_-tag in pET28b(+) bacterial expression vector (kanamycin resistance) was kindly provided by Prof. Tapas K. Kundu. All NPM1 mutants except for T46R contained this tag.

Mutant forms of NPM1 containing point mutations T78E and S112E were derived via megaprimer PCR [1] from the wild-type NPM1 plasmid using the standard T7 forward primer with corresponding mutagenic reverse primer listed in Supplementary Table 1 in the first PCR and obtained megaprimer and standard T7 reverse primer in the second PCR. The resulting PCR products were cloned into pET28b(+) vector using *NcoI* / *XhoI* restriction endonuclease sites. The T46R mutant of NPM1 was derived from the wild-type NPM1 plasmid using primers NdeI_NPM1_forward, NPM1_T46R_reverse and NPM1_stop_Xho_reverse. The obtained construct was treated with *NdeI*/*XhoI* restriction endonucleases and then cloned into pET28b(+) pretreated with the same endonucleases. NPM1-T46R protein form contained the N-terminal His_6_-tag cleavable by 3C protease, which was initially present in the vector used for cloning. After 3C cleavage the construct contained extra GPH residues at the N terminus.

For site-specific translational incorporation of the non-hydrolyzable analog of phosphoserine - PMA (p*S) - at positions 48 or 88 of NPM1 using amber codon suppression in the autonomous *E. coli* system [2], we created 48TAG and 88TAG variants. For NPM1-S88TAG we used the megaprimer method [1], NdeI_NPM1_forward primer, NPM1_S88_reverse mutagenic primer, His_HindIII_reverse primer and wild-type NPM1 template. The resulting PCR product was treated with *NdeI*/*HindIII* restriction endonucleases and then cloned into a pRBC vector (ampicillin resistance) [3] pretreated with the same endonucleases. For NPM1-S48TAG we used the wild-type NPM1 template in pRBC plasmid (this construct was obtained unintentionally when no desired mutations were introduced during previous cloning of NPM1-S48TAG via megaprimer PCR). Primers NdeI_NPM1_forward and HindIII_S43_reverse were used for introducing *HindIII* restriction site into the first PCR product. Second PCR fragment carrying *HindIII* site and desired mutation S48TAG was produced with primers HindIII_S43_S48_forward and His_HindIII_reverse. The first PCR product was treated with *NdeI/HindIII* endonucleases, the second – with HindIII/XhoI and then cloned into a pRBC-NPM1 vector pretreated with *NdeI*/*XhoI* endonucleases. The C-terminal His_6_-tag ensured facile removal of truncated and untagged products forming due to the presence in the *E. coli* cells of the Release Factor 1 responsible for translation termination at TAG codons [4].

The full-length human 14-3-3 protein isoforms beta (β), gamma (γ), eta (η), tau (τ), zeta (ζ), epsilon (ε), and sigma (σ) fused to the C terminus of the maltose-binding protein (MBP) were kindly shared by the laboratory of Prof. Gilles Travé (Université de Strasbourg, Illkirch, France) and are described in detail in our previous work [5]. All constructs were verified by DNA sequencing.

The cDNA of λ-phosphatase (Uniprot ID P03772, residues 1-221) was synthesized in Cloning Facility (https://cloning.tech/) as a construct containing an N-terminal His_6_-tag cleavable by human rhinovirus 3C protease for easy purification.

The SUMO-specific protease from *Brachypodium distachyon* with His_6_-tag (his_6_-bdSENP) in pET28b vector [2].

#### Protein expression and purification

Wild-type form of NPM1 and mutant forms T78E, S112E and T46R were expressed in *E. coli* BL21(DE3) cells in LB medium by the addition of IPTG up to 0.4-1 mM for 20 h at 30 °C after induction. For in-cell phosphorylation, different forms of NPM1 were co-expressed with the catalytic subunit of mouse PKA as described previously [6,7]. PKA was cloned into a low-copy pACYC vector (chloramphenicol resistance), which ensured that the target protein was expressed in excess over kinase [6]. The target NPM1 plasmid was transformed into competent *E. coli* BL21(DE3) cells already harboring the pACYC-PKA plasmid. Cells were grown in LB medium to an OD_600_ reading of 0.3-0.8 before inducing with 0.5-1 mM of IPTG. After induction, cultivation was continued for 48 h at 28-30 °C.

NPM1 forms containing p*S were expressed in BL21(DE3)ΔserC cells (the derivative of BL21(DE3) strain with the knocked out gene for phosphoserine aminotransferase (SerC)); Addgene accession number 197656) co-transformed simultaneously by three plasmids: pCDF-Frb-v1.0 (spectinomycin resistance; Addgene accession number 201923) and pSF-nhpSer (kanamycin resistance), where enzymes for p*S synthesis, the engineered tRNA and necessary factors are encoded (Supplementary Fig. 3), along with the appropriate pRBC plasmid encoding the corresponding NPM1 form with either p*S48 or p*S88, as described in detail earlier [2]. Expressions were performed as previously described [8]. Briefly, fresh transformations were prepared for every expression. The cells were recovered for 90 min in SOC media at 37 °C and plated onto LB/agar plates containing 50 μg/mL ampicillin, 30 μg/mL kanamycin and 50 μg/ml spectinomycin. After overnight growth at 37 °C, cells were scraped and used to inoculate 2YT media supplemented with 0.5% glycerol and the same antibiotics. These cultures were grown for 3-5 hours at 37 °C, at which point they were used to inoculate Terrific Broth expression cultures (12 g/L tryptone, 24 g/L yeast extract, 0.5% glycerol, 72 mM K_2_HPO_4_, 17 mM KH_2_PO_4_). Expressions were initiated by the addition of 0.4-0.5 mM IPTG once cells reached an OD_600_ ∼1.0. Cultures were harvested after 20–24 h of expression at 20-30 °C.

The cells with overexpressed proteins were harvested by centrifugation and resuspended in 20 mM Tris-HCl buffer pH 8.0 containing 200 mM NaCl, 10 mM imidazole, 0.01 mM phenylmethylsulfonyl fluoride. NPM1 forms with C-terminal His_6_-tag were purified using immobilized metal-affinity chromatography (IMAC) and gel-filtration. NPM1-T46R carrying cleavable N-terminal His_6_-tag was purified similarly with addition of subtractive immobilized metal-affinity chromatography (IMAC). Phosphorylated proteins were purified using the same protocol, while the His_6_-tagged PKA was efficiently removed. For NPM1 wt coexpressed with PKA gel-filtration step was replaced by chromatography on heparin sepharose.

Untagged full-length human 14-3-3γ was expressed in *E.coli* BL21(DE3) cells by IPTG induction (final concentration of 1 mM) and purified by ammonium sulfate fractionation, anion-exchange and size-exclusion chromatography as described earlier [7]. His-tagged full-length human 14-3-3ε was obtained as described earlier [7]. Expression of the MBP-14-3-3 constructs was the same, but these proteins were purified using a combination of MBPTrap column (Cytiva) and size-exclusion chromatography on Superdex 200 26/600 (GE Healthcare) [3,5]. 40-kDa MBP was obtained as a side product of MBP-14-3-3 expression and purification upon gel-filtration [3]. His_6_-PKA was obtained by IMAC and gel-filtration as described earlier [7]. λ-phosphatase was purified by IMAC1, 3C proteolysis, IMAC2 and gel-filtration in this work. His_6_-bdSENP protease was purified by IMAC and dialyzed against SEC buffer.

Protein concentrations were determined on a NP80 nanophotometer (Implen, Munich, Germany) at 280 nm using the extinction coefficients listed in Supplementary Table 2.

#### Recombinant NPM1 phosphopeptides

To study 14-3-3 binding to internal NPM1 phospho-motifs, we obtained recombinant NPM1 peptides containing p*S instead of Ser48, Ser106 or Ser260 (Fig. 5B). To produce these peptides in *E. coli* cells, a special fusion construct was developed on the basis of the pRBC plasmid, which contained the following parts: an N-terminal His_6_-tag, small ubiquitin-related modifier (SUMO) domain from *Brachypodium distachyon* (bdSUMO), the corresponding peptide sequence with an additional tryptophan residue on its C-terminal end for easier peptide detection upon purification, a thrombin cleavage site (leaving extra LVPR residues at the peptide’s C terminus) and enhanced yellow fluorescent protein (YFP) (Fig. 5B). A forward primer for PCR included *BamHI* site and the whole sequence of a peptide, where the Ser48 (or Ser106 or Ser260) codon was replaced by a TAG codon (see primer sequences in Supplementary Table 1). We used ‘M13_forward’ as a reverse primer. The resulting PCR fragment was treated with *BamHI*/*XhoI* endonucleases and cloned into the pRBC plasmid.

Plasmids pCDF-Frb-v1.0, pSF-nhpSer and pRBC-His_6_-bdSUMO-peptide-YFP required for autonomous p*S-peptide biosynthesis (Supplementary Fig. 3 [2]) were cotransformed simultaneously into BL21(DE3)ΔserC cells. Fresh transformations were performed for every expression. The cells were recovered for 90 min in SOC media at 37 °C, and plated onto autoinducing 4YT/agar plates containing 0.4% lactose, 0.1% glucose, 1.2% glycerol, 50 g/mL ampicillin, 30 g/mL kanamycin and 50 g/ml spectinomycin. After overnight growth at 37 °C and several days at room temperature, each colony was resuspended in water or medium and then protein expression was evaluated by measuring YFP fluorescence using Clariostar plus microplate reader (BMG Labtech, Offenburg, Germany) with a set of bandpass filters (482 ± 16 nm and 530 ± 40 nm). Best expressing colonies were scraped and used to inoculate 2YT media supplemented with 0.5% glycerol and antibiotics. These cultures were grown for 3-5 hours at 37 °C, at which point they were used to inoculate Terrific Broth expression cultures (12 g/L tryptone, 24 g/L yeast extract, 0.5% glycerol, 72 mM K_2_HPO_4_, 17 mM KH_2_PO_4_). Expressions were initiated by the addition of 0.5 mM IPTG once cells reached an OD_600_ ∼1.5-2.0. Cultures were harvested after 20–24 h of expression at 30 °C.

In addition, we produced a recombinant C-terminal NPM1 peptide containing Ser293. This peptide has an intact NPM1’s C-terminus and therefore the design of the above described constructs with extra LVPR residues after thrombin cleavage was not suitable. For this case, we used primers DuetUP2 and NPM_pept_S293_R to modify the construct and obtain a new one, having only an N-terminal His_6_-bdSUMO fusion tag followed by the peptide sequence, which could be optionally phosphorylated by PKA during co-expression (Fig. 5B). The resulting PCR product was cloned into a pET28b(+) vector treated with *NcoI*/*XhoI* endonucleases. The plasmid was transformed into BL21(DE3)-pACYC-PKA cells so that the protein was phosphorylated during co-expression with PKA (LB media, inducing with 0.5 mM of IPTG when OD_600_ = 0.6, cultivation for 48 h at 30 °C after induction). To obtain the same peptide in the unphosphorylated form, His_6_-bdSUMO-S293 peptide was expressed in cells BL21(DE3) without PKA.

The cells with overexpressed proteins were harvested by centrifugation and resuspended in 20 mM Tris-HCl buffer pH 8.0 containing 200 mM NaCl, 10 mM imidazole, 0.01 mM phenylmethylsulfonyl fluoride (PMSF). The purification protocol for p*S peptides consisted of the following steps: IMAC, specific cleavage of His_6_-bdSUMO by His_6_-bdSENP, subtractive IMAC, gel-filtration (HiLoad 16/600 Superdex 75 (GE Healthcare)), FITC-labeling, specific cleavage of YFP by thrombin, separation from unreacted FITC using HiTrap Desalting column (GE Healthcare), precipitation of remaining protein in 50% acetonitrile and gel-filtration on Superdex Peptide 10/300 column (Amersham Pharmacia) in 30% acetonitrile (ACN) with 0.1% trifluoroacetic acid (TFA). For the peptide with pS293 (or S293) the protocol was modified to minimize unspecific hydrolysis and dephosphorylation: the lysis buffer contained EDTA-free inhibitors of proteases (PierceTM Protease Inhibitor Mini Tablets, cat. no. A32955) and phosphatase inhibitor tablets (PhosSTOP™, cat. no. 4906845001), IMAC step 1 [HisCap 6FF column (Smart-Lifesciences); A: 50 mM sodium phosphate, pH 8.0, 425 mM NaCl; B: 50 mM sodium phosphate, pH 8.0, 425 mM NaCl, 500 mM imidazole], gel-filtration on Superdex 75 16/600, removal of His_6_-bdSUMO by His_6_-bdSENP, IMAC step 2, precipitation of the remaining protein in 30% ACN and clarification by centrifugation, gel-filtration on Superdex 30 16/600 (Cytiva) in 30% ACN/0.1% TFA, FITC labeling, separation from unreacted FITC using Sephadex G10 resin. The addition of tryptophan to the sequence of peptides allowed us to analyze different purification steps using SDS-PAGE with 2,2,2-Trichloroethanol (TCE, Sigma) [9] according to the Stain-free technology® implemented in Bio-Rad Gel-Doc EZ system, immediately after the gel run.

#### Far-UV circular dichroism spectroscopy

The secondary structure of wild-type NPM1 and its glutamate mutants T78E and S112E was analyzed by circular dichroism (CD). Protein samples (0.8–1 mg/mL) were dialyzed overnight against 20 mM Tris (pH 7.5) containing 150 mM NaCl and 5 mM β-mercaptoethanol (βME) and then centrifuged for 10 min at 4 °C and 20,000 g. Protein concentration in supernatants was determined spectrophotometrically at 280 nm and then far-UV (190-280 nm) CD spectra of the samples were recorded at RT. Spectra were recorded twice at a rate of 0.4 nm/s with 1 nm steps in 0.1 cm quartz cuvette on a Chirascan Circular Dichroism Spectrometer (Applied Photophysics) equipped with temperature controller, and then the signal from buffer, filtered through 0.22 µm membrane, was subtracted. Secondary structure elements were estimated using DichroWeb server [10] by CDSSTR algorithm with a set 4 (best fit for the wild-type NPM1) and 7 of reference proteins (best fits for the mutants of NPM1) and compared with the expected secondary structure content within NPM1 having the OD and C-terminal domains folded (i.e., ∼17% α-helices, ∼37% β-strands, ∼46% unstructured regions).

#### In vitro phosphorylation/dephosphorylation and gel-electrophoresis

For phosphorylation reaction the protein (1–1.7 mg/ml) was incubated at 37 °C in the buffer containing 12.5 mM HEPES pH 7.3, 2.5 mM NaCl, 1 mM MgCl_2_, 0.3 mM ATP with addition of 5 mM phosphate buffer (to inhibit dephosphorylation) in the presence or in the absence of the catalytic subunit of recombinant mouse PKA (∼60 µg/ml). The reaction mixture for dephosphorylation contained ∼60-90 µM of phosphorylated protein in 20 mM Tris-HCl pH 7.8 with 150 mM NaCl, 1.9 mM MnCl_2_, 5 mM DTT. The reaction was started by the addition of PKA or λ-phosphatase, respectively. The shift of the electrophoretic mobility due to protein phosphorylation/dephosphorylation was analyzed on native PAGE and SDS-PAGE gels as described earlier [6,11]. The small heat shock protein HSPB6 was used as a reference PKA substrate [12].

#### Size-exclusion chromatography

Oligomeric state of NPM1 was examined by loading its preparations on a Superdex 200 Increase 5/150 column (GE Healthcare) equilibrated by 20 mM TRIS-HCl pH 7.6 buffer containing 150 mM NaCl and 3 mM βME (SEC buffer) and operated at a 0.45 ml/min flow rate. The elution profiles were followed by absorbance at 280 nm and then normalized by peak maxima when necessary. To assess the apparent Mw for the peak maxima of the proteins in question we used column calibration with BSA trimer (198 kDa), BSA dimer (132 kDa), BSA monomer (66 kDa) and α-lactalbumin (15 kDa).

Interactions of NPM1 variants with 14-3-3 were studied by loading on a Superdex 200 Increase 10/300 column (GE Healthcare) prepared as above (flow rate of 0.8 ml/min) of individual proteins or their mixtures preincubated in SEC buffer for at least 20 min at room temperature. Protein concentrations are indicated in figures. During the elution, fractions were collected and their protein content was analyzed by SDS-PAGE. Additionally, we subjected 14-3-3/NPM1 mixtures to native PAGE in the glycine-TRIS system [13]. All SEC experiments were performed at least three times and the most typical results are presented.

#### SEC-MALS

Absolute Mw for NPM1 variants was determined by size-exclusion chromatography coupled with multi-angle light scattering (SEC-MALS). To this end, a Superdex 200 Increase 10/300 column operated at flow rate of 0.8 ml/min by a ProStar 210/335 chromatography system (Varian Inc., Melbourne, Australia) was connected to a 335 UV/Vis diode-array detector and a miniDAWN multi-angle laser light scattering detector (Wyatt Technology). The miniDAWN detector was calibrated relative to the scattering from toluene and, together with concentration data obtained from the UV detector at 280 nm, was exploited for determining the Mw distribution of the eluted protein species. Data processing was performed in ASTRA 8.0 software (Wyatt Technologies) considering dn/dc equal to 0.185 and using extinction coefficients listed in Supplementary Table 2.

#### FITC labeling of recombinant NPM1 peptides

Peptide labeling was achieved in 0.1 M carbonate buffer, pH 8.5-9.2, by mixing 3.5-20 mg/ml solution of the peptide-YFP (or ∼3 mg/ml solution of the peptide in the case of pS293) with < 1/10 volume of the FITC stock solution in DMSO (maintaining peptide:FITC molar ratio equal to 1:5-1:8). The mixtures were wrapped with aluminum foil and incubated overnight at room temperature.

The FITC-labeled peptide-YFP constructs were separated from the unreacted label by chromatography on a Sephadex-G25 desalting column (5 ml, GE Healthcare) equilibrated using a 20 mM Tris-HCl buffer, pH 7.5, 150 mM NaCl. After that, peptide-YFP constructs were specifically proteolysed by thrombin protease (at room temperature, ON) and precipitated in 50% acetonitrile. Final purification of the peptides was performed via SEC on a 24 ml Superdex Peptide 10/300 column (GE Healthcare) in 30% ACN/0.1% TFA at a 0.5 ml/min flow rate. The runs were operated by the Prostar 335/363 system (Varian Inc., Australia) enabling continuous detection of full-spectrum absorbance and of FITC fluorescence at λ_ex_ = 480 nm and λ_em_ = 520 nm. Elutions were analyzed using SDS-PAGE with TCE and evaporized using rotational vacuum concentrator RVC 2-25 CDplus (Martin Christ Gefriertrocknungsanlagen GmbH, Germany) at 25 °С 1200 rpm for 4 h.

The FITC-labeled peptides pS293 and S293 were separated from the unreacted label by SEC on a 1 ml Sephadex G10 column equilibrated and run using a 20 mM Tris-HCl buffer, pH 7.5, 150 mM NaCl at a 0.3 ml/min flow rate. The runs were operated by the Prostar 335 system (Varian Inc., Australia) following the full-spectrum absorbance.

The resulting spectrum, and absorbance at 280 (A_280_) and 493 nm (A_493_) in particular, was used to estimate the phosphopeptide concentration (C_peptide_), taking into account FITC absorbance at 280 nm using the formula [14]:

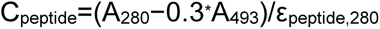

where ε_peptide, 280_ is the extinction coefficient of the peptide at 280 nm (for all p*S peptides in the study – 5500 M^−1^ cm^−1^, for pS293 peptide – 11,000 M^−1^ cm^−1^). FITC concentration was determined from the FITC-specific absorbance at 493 nm using the extinction coefficient of 68,000 M^−1^ cm^−1^. The sequence of peptides and the completion of the labeling reaction were confirmed by LC-MS on a Bruker Impact II mass-spectrometer (Supplementary Fig. 7). The labeling efficiency was up to 80% for all phosphopeptides. Labeled peptides were stored in aliquots at −80 °C.

#### Fluorescence anisotropy

Fluorescence anisotropy was measured at 25 °C on a Clariostar plus microplate reader (BMG Labtech, Offenburg, Germany) using a set of bandpass filters (482 ± 16 nm and 530 ± 40 nm), FITC labeled phosphopeptides (400 nM stock concentration, on a 20 mM HEPES-NaOH buffer, pH 7.5, 150 mM NaCl, 0.08% Tween 20) and 384-well plates (Black Nunc™ 384-Shallow Well Standard Height Polypropylene Storage Microplates, Thermofisher scientific, cat. no. 267460). A series of concentrations of MBP-14-3-3 proteins in a 20 mM HEPES-NaOH buffer, pH 7.5, 150 mM NaCl (105 μl each) was diluted by a 400 nM solution of either FITC-labeled peptide to obtain 120 μl samples containing 50 nM peptide, MBP-14-3-3 (from 0 to 120 μM) and 0.01% Tween 20. Then, each sample was split into three 30-μl aliquots, the plates were centrifuged for 3 min at 25 °C at 150 g to remove air bubbles, and the fluorescence anisotropy data were recorded. The triplicate measurements were used for the data presentation and global fitting. Titration of the FITC-phosphopeptide by lysozyme (0-80 μM) or MBP (0-80 μM) and titration of free FITC by MBP-14-3-3η were performed as controls. To determine *K*_D_ values, the binding curves were approximated using the quadratic equation [15] in Origin 9.0 (OriginLab Corporation, Northampton, MA, USA):

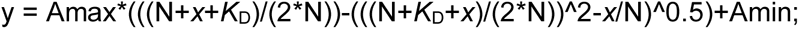

where Amax is fluorescence anisotropy (mA) at saturation, Amin - fluorescence anisotropy (mA) at 0 μM protein, N – concentration of NPM1 peptide (μM), *K*_D_ – dissociation constant (μM), and y – fluorescence anisotropy (mA) at *x* protein concentration.

To exclude the effect of FITC-labeling efficiency on the quality of the fitting and affinity constants determined, we varied the fixed amount of titrated FITC-peptide in the range 10-50 nM and performed fitting. Given the significantly exceeding micromolar concentrations of titrating 14-3-3, the exact nanomolar peptide concentration and the exact labeling efficiency did not matter for the fitting whatsoever.

Another set of experiments involved the addition of an ethanolic stock solution of FSC (final concentration 100 μM) to investigate the influence of this compound on stabilization (or inhibition) of 14-3-3 complexes with NPM1 peptides. The binding curves obtained in the presence of FSC were compared with the binding curves obtained in the presence of the equivalent amount of ethanol added to the sample instead of FSC (the final concentration of ethanol did not exceed 1.5%).

#### Identification of NPM1 phosphosites using LC-MS analysis

NPM1 digestion was performed using Trypsin Gold (Promega, USA) according to the manufacturer’s protocol. UPLC-MS analysis was performed on Impact II QTOF high-resolution mass-spectrometer (Bruker Daltonik, Germany) equipped with Apollo II ESI ion source (Bruker Daltonik, Germany) coupled to Elute UPLC (Bruker Daltonik, Germany) on Waters Acquity HSS T3 1.8 µm 2.1 × 100 mm reverse phase column (Waters, Ireland) with the following conditions: 50 µL injection volume, gradient elution at 0.25 mL/min from 5% to 60% B in 30 min, then to 95% B in 3 min (A: 0.1% formic acid in water, B: 0.1% formic acid in acetonitrile), column at 30 °C, post-column 1:20 flow split, ion source in positive mode, HV capillary at 4.5 kV, spray gas – nitrogen at 1.0 bar, dry gas – nitrogen at 5.0 L/min 200 °C, full spectra scan range m/z 100-2200 at 2 Hz scan rate, data-dependent MS/MS spectra acquisition via CID at a dynamic rate of 2–6 Hz, cycle time 2 sec, preferred charge state 2–6, nitrogen as collision gas at variable collision energy (23 eV at m/z 300, 65 eV at m/z 1300), automatic internal calibration with sodium trifluoroacetate solution. Spectra processing and protein identification were performed in BioPharma Compass 3.1.1 (Bruker Daltonik, Germany) and Mascot 2.8.1 (Matrix Science, United Kingdom). Recombinant NPM1 peptides p*S48, p*S106, p*S260 and pS293 were analyzed by LC-MS nearly identically, with the caveat that no proteolytic digestion took place.

## Supplementary Tables

**Table S1.**
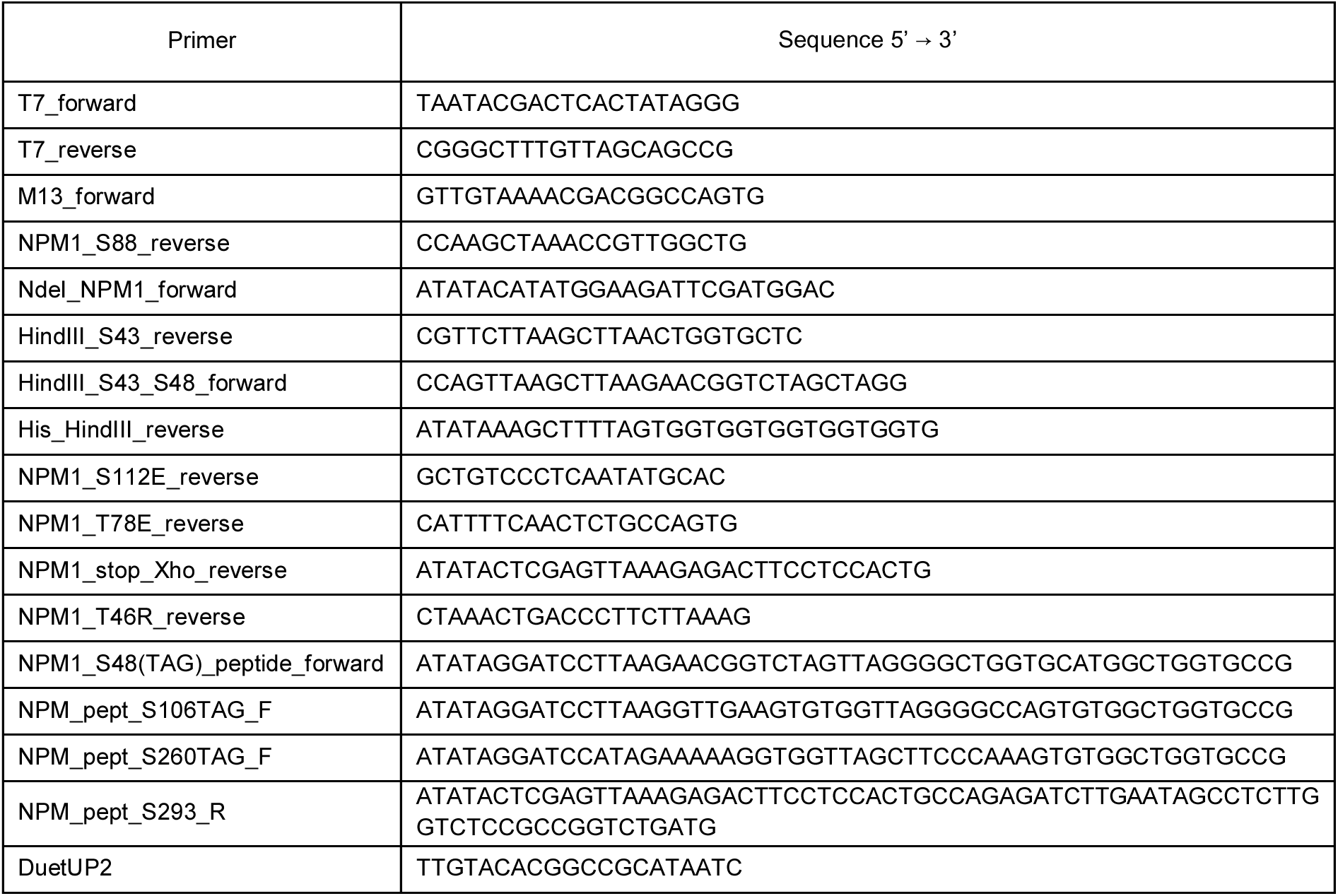
Primer sequences used in this study.

**Table S2.** Properties of the proteins studied.

| Protein | MW, Da | pI | Molar ext. coef.<br>M<sup>-1</sup> cm<sup>-1</sup> | Ext. coef.<br>(mg/mL)<sup>-1</sup> cm<sup>-1</sup> |
| --- | --- | --- | --- | --- |
| MBP-14-3-3ε | 69505.8 | 4.80 | 95230 | 1.370 |
| MBP-14-3-3γ | 68634.5 | 4.90 | 98210 | 1.431 |
| MBP-14-3-3ζ | 68077.0 | 4.90 | 93740 | 1.377 |
| MBP-14-3-3τ | 68298.4 | 4.90 | 93740 | 1.373 |
| MBP-14-3-3η | 68752.9 | 4.90 | 95230 | 1.385 |
| MBP-14-3-3σ | 68308.2 | 4.80 | 92250 | 1.350 |
| MBP-14-3-3β | 68616.5 | 4.90 | 93740 | 1.366 |
| 14-3-3γ | 28302.6 | 4.80 | 31860 | 1.130 |
| his-14-3-3ε | 32548.0 | 4.85 | 30370 | 0.930 |
| NPM1 wt | 33639.9 | 4.81 | 16960 | 0.504 |
| NPM1-T78E | 33667.9 | 4.79 | 16960 | 0.504 |
| NPM1-S112E | 33681.9 | 4.79 | 16960 | 0.504 |
| NPM1-T46R<sup>a</sup> | 32921.2 | 4.70 | 16960 | 0.515 |
| NPM1-p*S88 | 33717.9 | 4.81 | 16960 | 0.504 |
| NPM1-p*S48 | 33717.9 | 4.81 | 16960 | 0.504 |
| his-bdSUMO-peptide-p*S48-YFP | 40582.5 | 5.86 | 30370 | 0.748 |
| peptide-p*S48-YFP | 28800.5 | 5.85 | 28880 | 1.003 |
| his-bdSUMO-peptide-p*S106-YFP | 40667.7 | 5.93 | 30370 | 0.747 |
| peptide-p*S106-YFP | 28885.7 | 5.99 | 28880 | 1.000 |
| his-bdSUMO-peptide-p*S260-YFP | 40665.6 | 5.86 | 30370 | 0.747 |
| peptide-p*S260-YFP | 28883.6 | 5.85 | 28880 | 1.000 |
| his-bdSUMO-peptide-pS293 | 13698.1 | 5.70 | 12490 | 0.912 |
| MBP | 40136.3 | 5.04 | 66350 | 1.653 |
| lysozyme | 14388.0 | 11.35 | 38940 | 27.21 |

<sup>a</sup> all NPM1 mutants except for T46R contained uncleavable C-terminal His<sub>6</sub>-tags, therefore their listed properties correspond to the tagged version. For the T46R protein, which contains the N-terminal His<sub>6</sub>-tag cleavable by 3C protease, the parameters are shown for the untagged version. Its tagged version has calculated Mw of 35,067.5 Da and pI of 4.89.

**Table S3.**
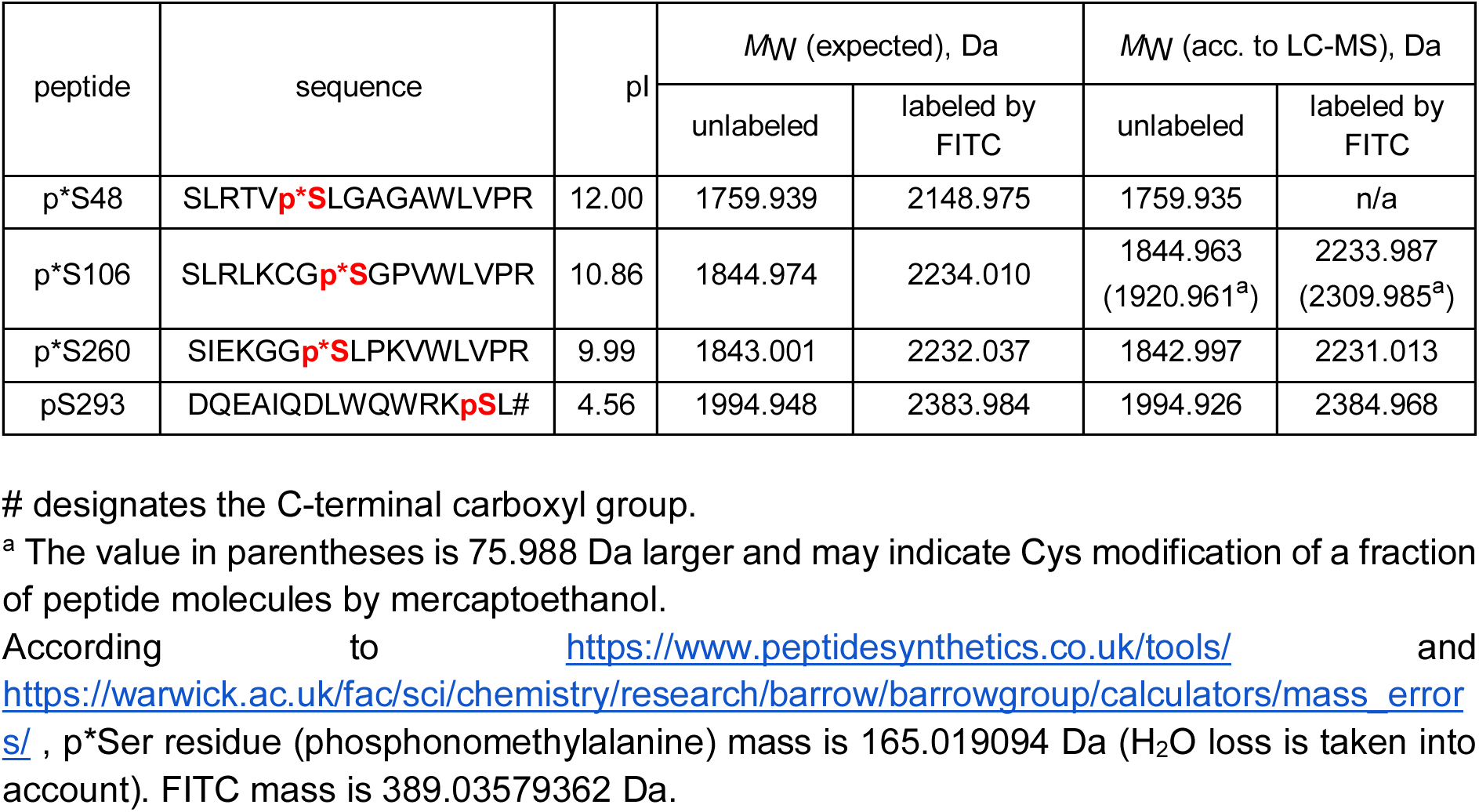
Monoisotopic masses of the peptides and their FITC-labeled derivatives.

## Supplementary Figures

**Fig. S1.**
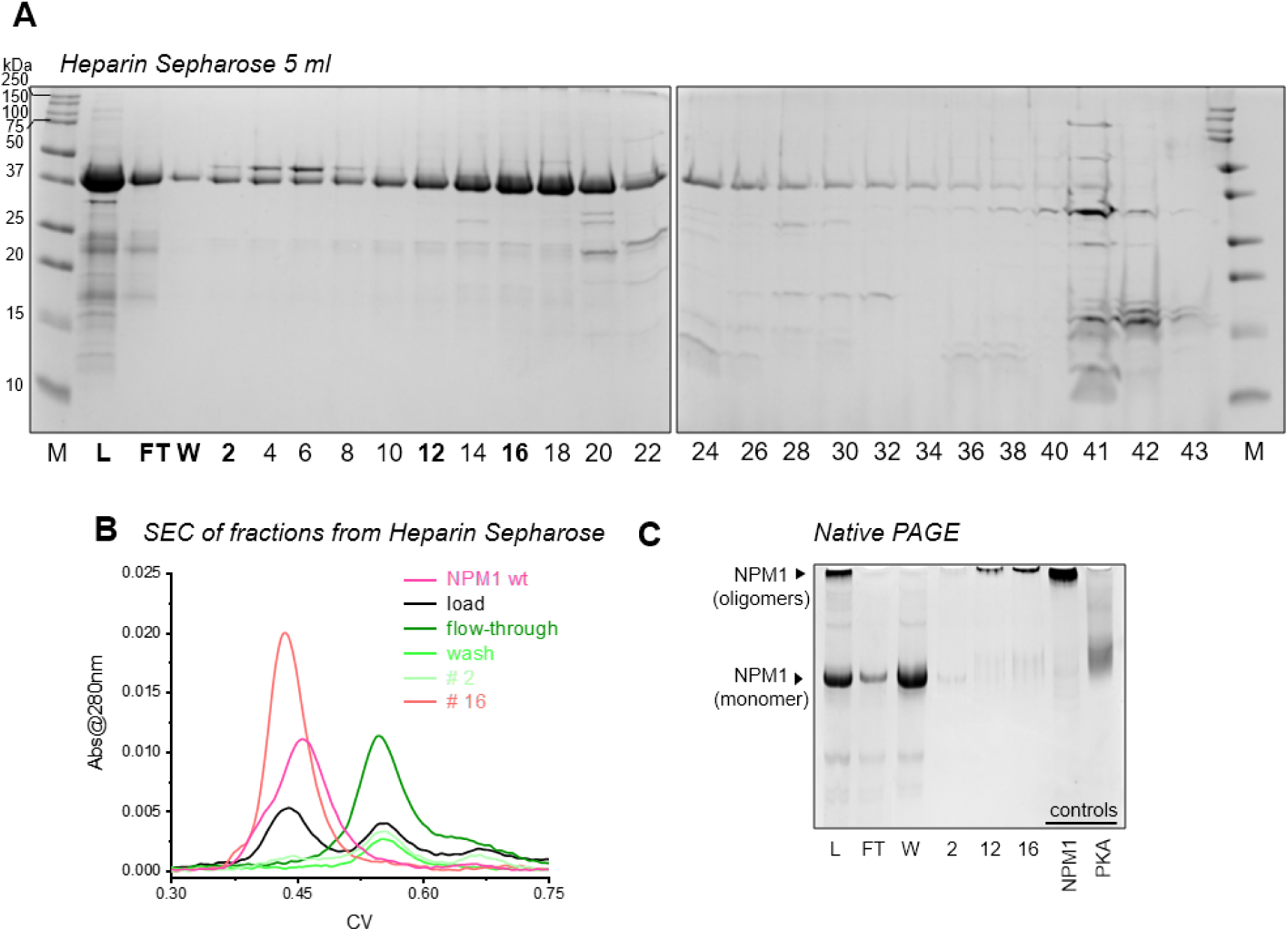
NPM1 phosphorylated by PKA in *E.coli* cells contains two oligomeric forms differing by affinity for heparin. A. Chromatography profile of phosphorylated NPM1 obtained from a Heparin Sepharose (GE Healthcare) column and analyzed by SDS-PAGE. M, standard proteins, L loading sample, FT - unbound fraction, W - wash with low salt buffer, numbers correspond to the fraction numbers obtained during gradient elution with increasing salt concentration. Bold fractions were analyzed by SEC on a Superdex 200 Increase 5/150 column (flow rate 0.45 ml/min, B) and native PAGE (C). Note that NPM1 incapable of heparin binding has high mobility on native PAGE and reduced size and hence likely corresponds to the phosphorylated and disassembled oligomer.

**Fig. S2.**
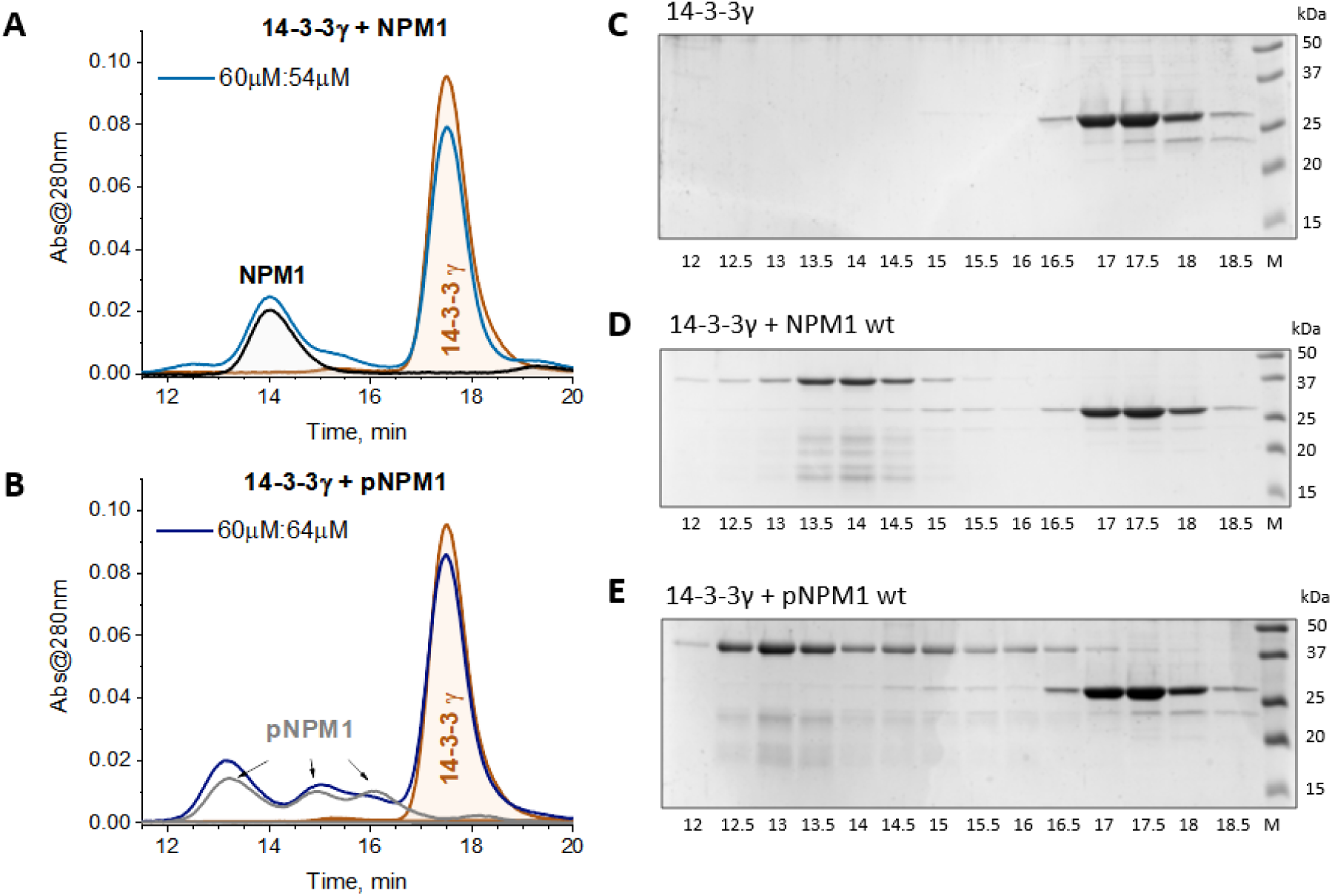
Interaction of unphosphorylated or PKA-phosphorylated wild-type NPM1 with 14-3-3γ. A and B. SEC profiles of individual 14-3-3γ, individual unphosphorylated NPM1 (A), individual PKA-phosphorylated NPM1 (B) or the 14-3-3/NPM1 mixtures obtained using a Superdex 200 Increase 10/300 column (flow rate 0.8 ml/min). C-E. SDS-PAGE analysis of the fractions obtained during the SEC runs of individual 14-3-3γ (C) or the 14-3-3 mixtures with either unphosphorylated NPM1 (D) or NPM1 phosphorylated by PKA (E). NPM1 phosphorylation was achieved by co-expressed PKA in *E.coli* cells. Lanes are labeled according to the elution times in min. M - protein ladder.

**Fig. S3.**
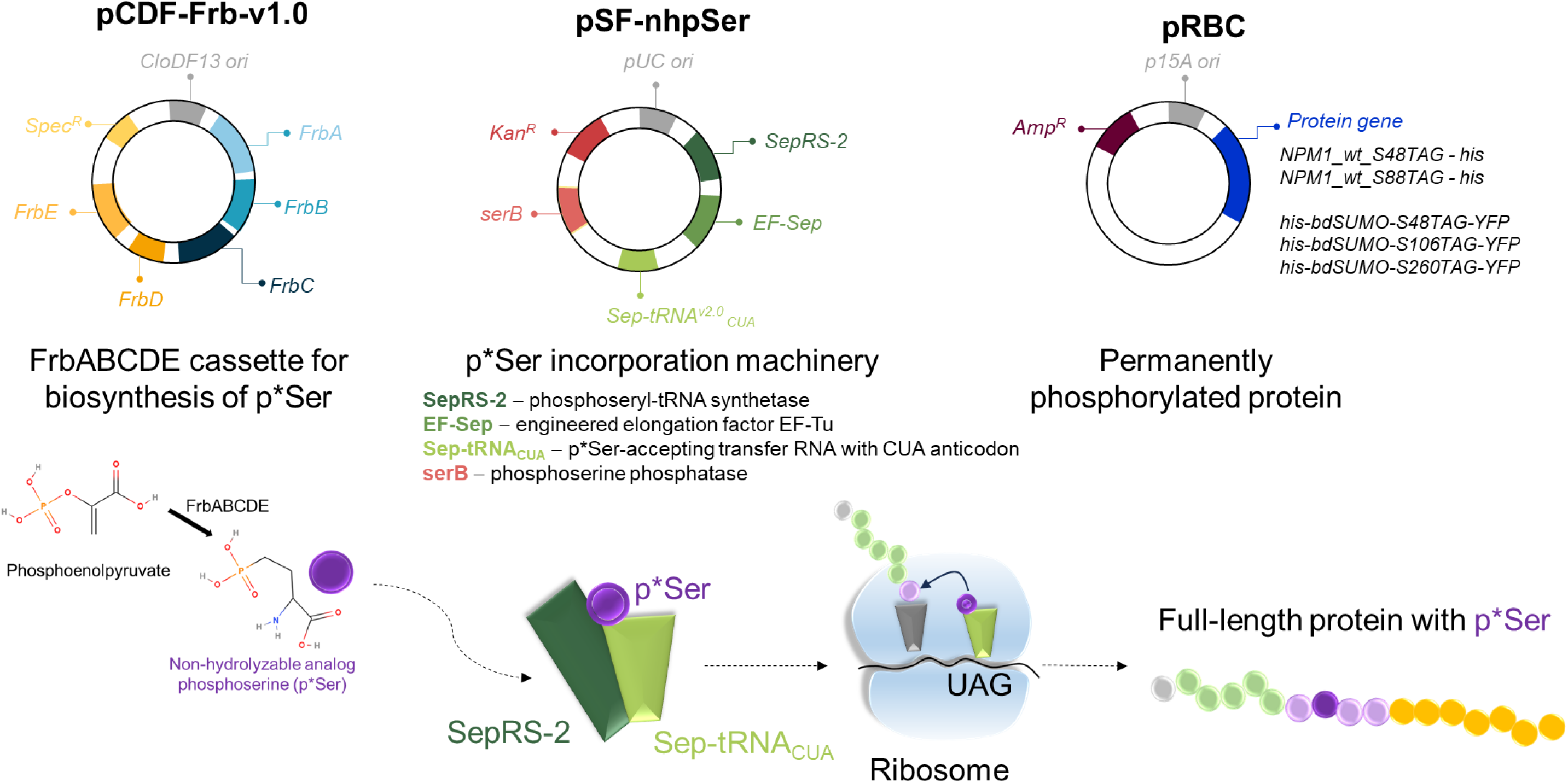
Schematic showing the three plasmids used for the biosynthesis of NPM1 peptides with translational incorporation of phosphonomethylalanine (p*S) autonomously produced from the natural metabolite phosphoenolpyruvate in *E. coli* [2]. p*S incorporation in place of the TAG codon was achieved in the BL21(DE3)ΔserC strain with the serC gene encoding phosphoserine aminotransferase knocked out to minimize the buildup of phosphoserine [2].

**Fig. S4.**
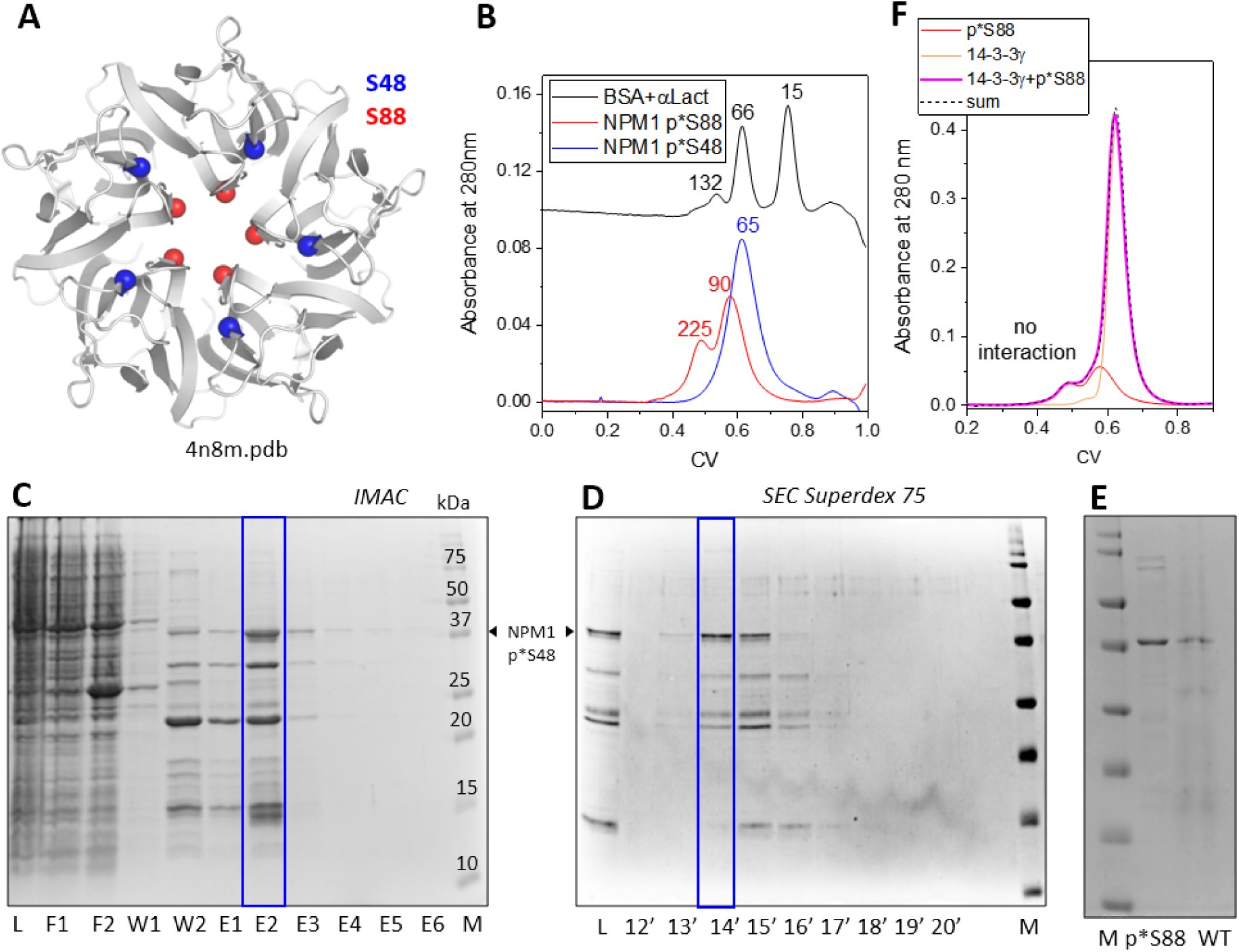
NPM1 singly modified by p*S at either 48th or 88th serine in the oligomerization domain. A. NPM1 structure showing the location of the serines targeted for PMA encoding. B. Analytical SEC profiles showing the effect of these single phosphorylation-related modifications on the elution of NPM1. The SEC profile for the standard proteins is shown for comparison. Apparent Mw values determined from column calibration are indicated above the peaks in kDa. Note that p*S48 leads to more complete oligomer disassembly than p*S88. C and D. SDS-PAGE analysis of the fractions obtained in the course of p*S48 NPM1 purification by IMAC (C) and SEC (D). Arrow shows the position of the p*S48 NPM1 variant. M - protein ladder (Mw values are indicated in kDa in panel C). E. SDS-PAGE analysis of the purified p*S88 NPM1 variant and the wild-type NPM1. F. SEC profiles showing the inability of p*S88 NPM1 variant to interact with 14-3-3γ.

**Fig. S5.**
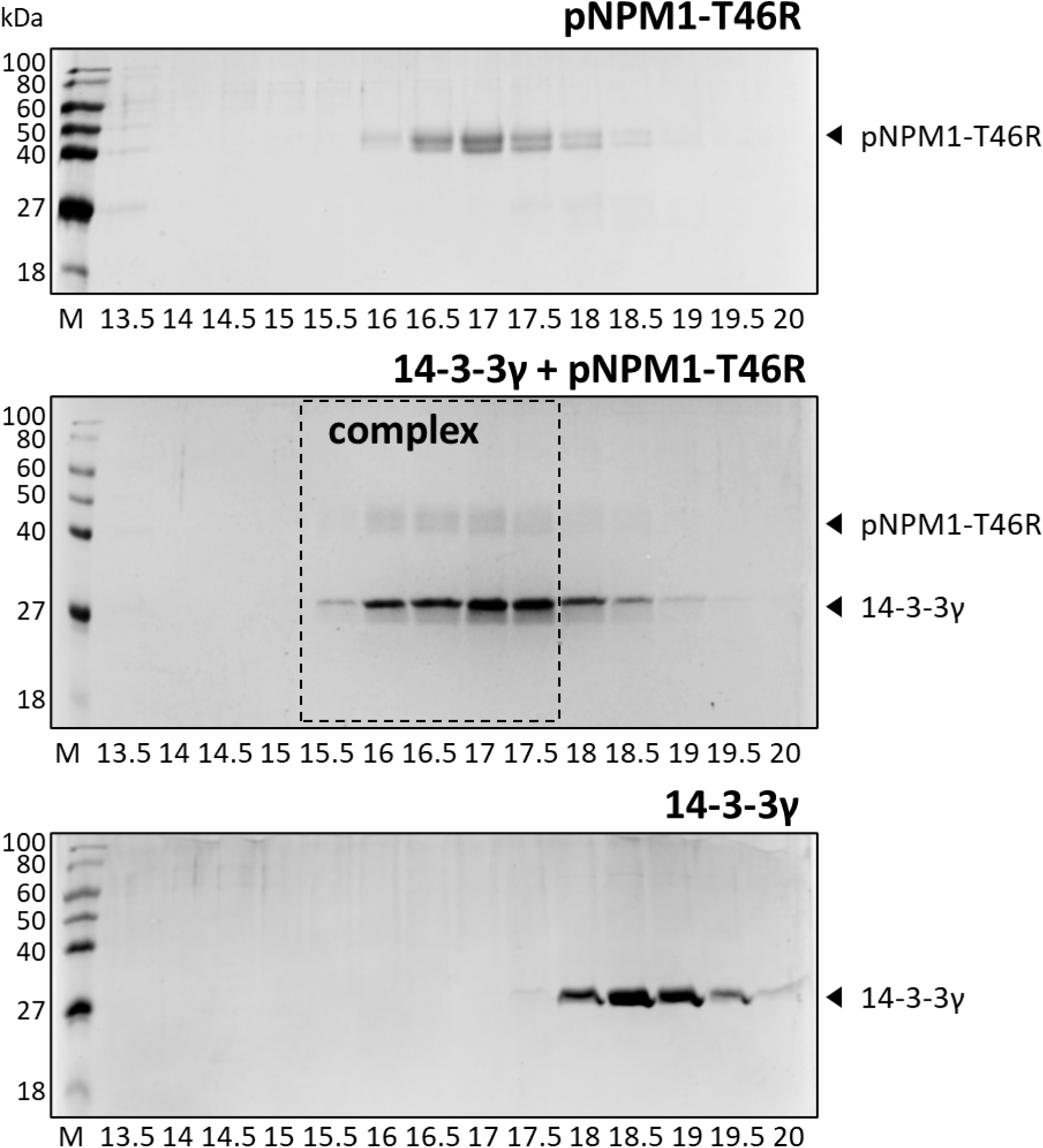
SDS-PAGE analysis of fractions obtained during the SEC runs of the individual phosphorylated T46R mutant of NPM1 (top), of its mixture with 14-3-3γ (middle), or of individual 14-3-3γ (bottom). The co-elution of the two proteins (dashed rectangle in the middle) indicates the formation of the protein-protein complex.

**Fig. S6.**
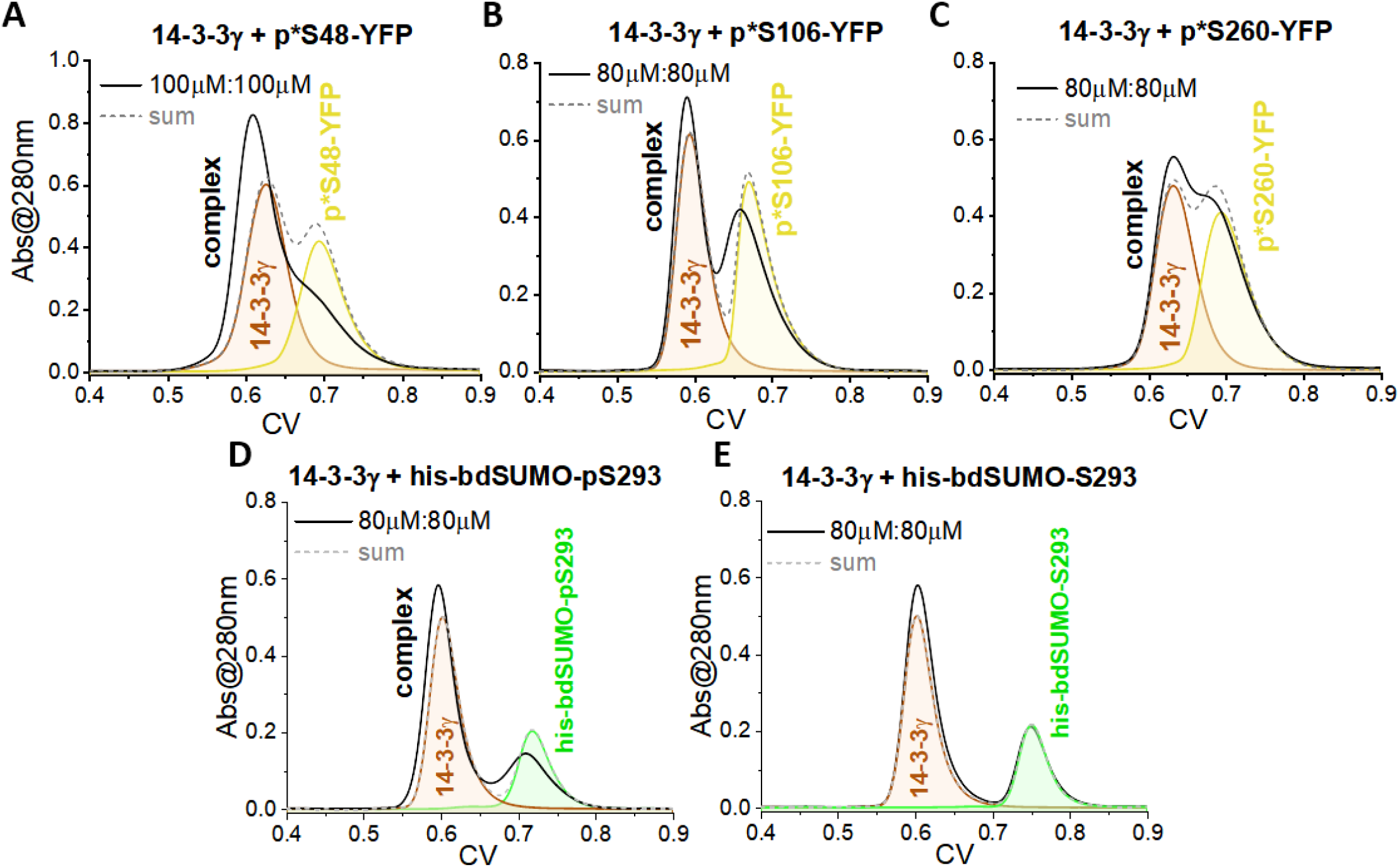
SEC analysis of the interaction between 14-3-3 and recombinant NPM1 peptides in a form of fusion proteins with either the C-terminal YFP (p*S peptides 48, 106, 260) or N-terminal his-bdSUMO (S293 peptides). Molar concentrations used for the analysis are indicated. Dashed lines correspond to the algebraic sums of the 14-3-3 and fusion protein profiles and are shown for reference. Individual profiles of 14-3-3 or fusion proteins are labeled. Note the different efficiencies of the complex formation for the different p*S and pS peptide fusions (A-D) and the absence of the interaction in the case of the unphosphorylated S293 peptide (E). Column: Superdex 200 Increase 5/150 column (flow rate 0.45 ml/min). pS293 peptide fusion phosphorylation was achieved in co-expression with PKA. p*S peptides contained the non-hydrolyzable pSer analog phosphonomethylalanine incorporated using the PermaPhos technology [2].

**Fig. S7.**
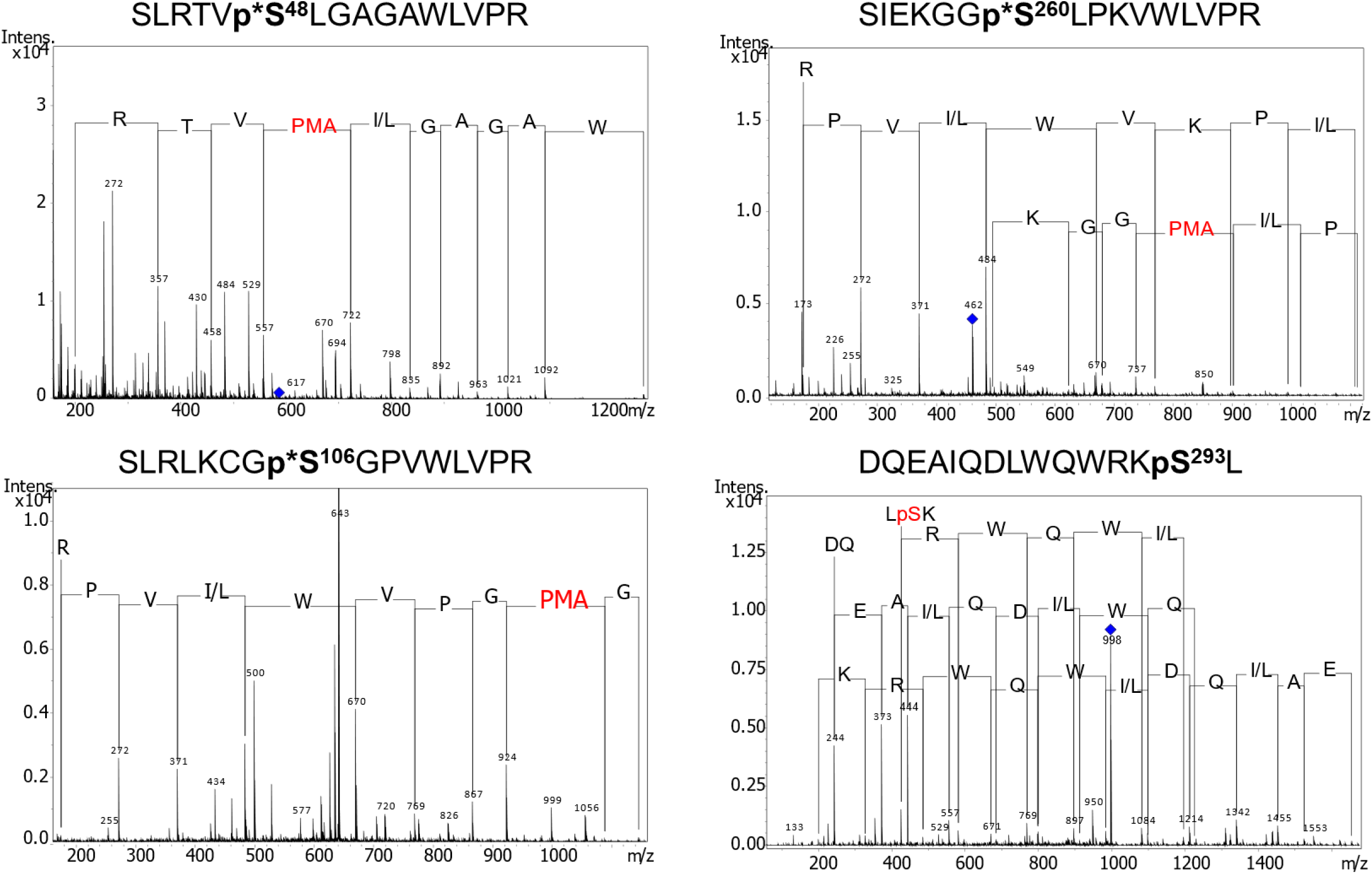
MS^2^ spectra demonstrating the identity of the studied recombinantly produced peptides containing either p*S (phosphonomethylalanine, PMA) or pS (phosphoserine).

**Fig. S8.**
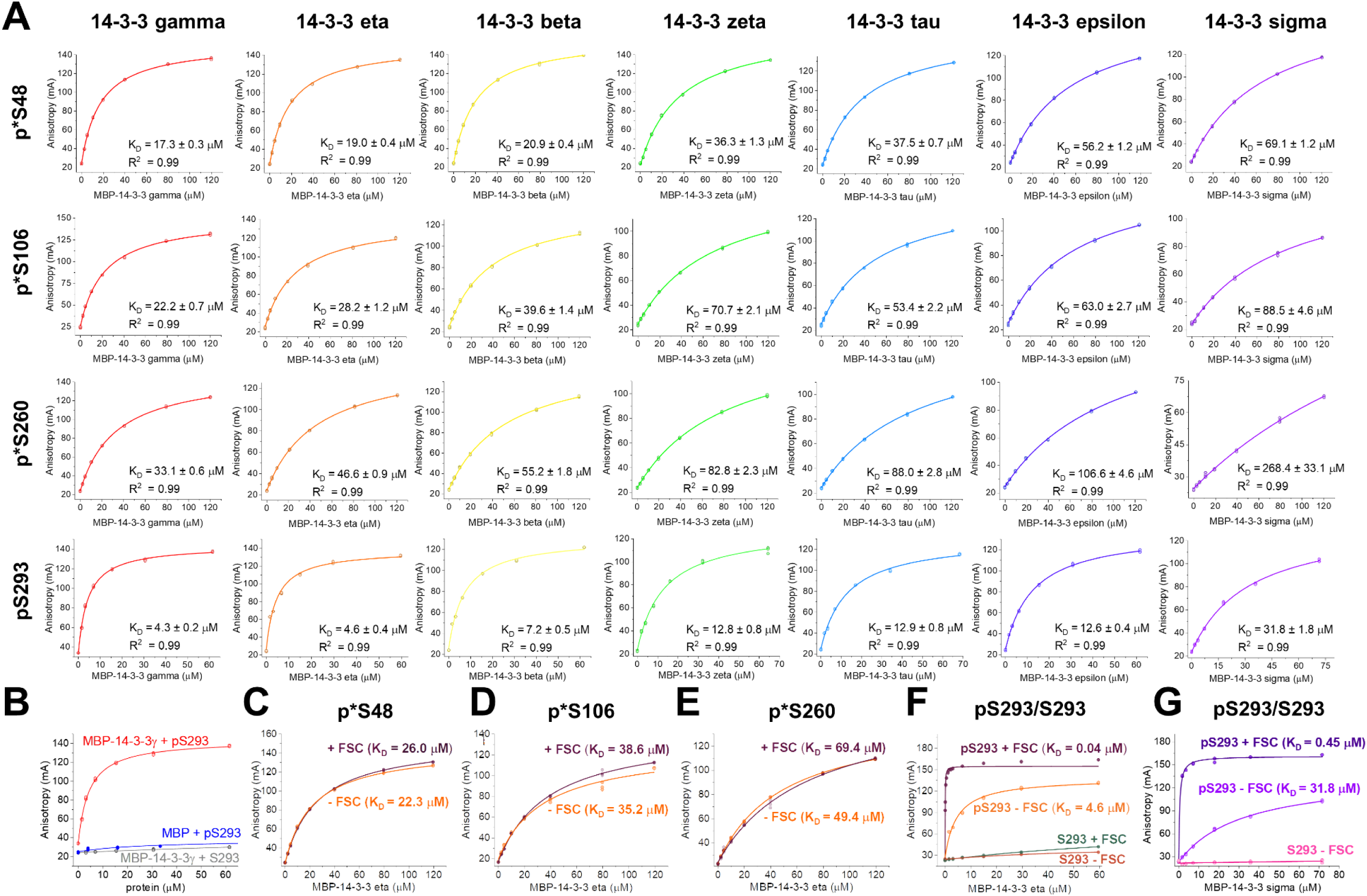
Titration of NPM1 phosphopeptides by the seven human 14-3-3 isoforms (A,B) and modulation of the interaction by fusicoccin (FSC) (C-G). Control experiments include titration of the FITC-labeled pS293 peptide by MBP (B), titration of the unphosphorylated FITC-labeled S293 peptide by MBP-14-3-3γ (B), titration of the unphosphorylated FITC-labeled S293 peptide by MBP-14-3-3η (F) or MBP-14-3-3σ (G), in the absence or presence of 100 μM FSC. Control titration of the phosphorylated FITC-labeled phosphopeptides by lysozyme or titration of free FITC by MBP-14-3-3 are shown in Fig. 5. Each titration was performed in triplicate and all data were used for fitting and presentation without correcting for the initial anisotropy values. The mean equilibrium dissociation constants (*K*_D_) are shown for each curve along with the standard deviation and the coefficient of determination *R*^2^. For titrations with FSC, the corresponding micromolar *K*_D_ values are indicated in parentheses (coefficient of determination *R*^2^ was equal to 0.99 and is not indicated on plots).

